# Odor-evoked activity outside canonical areas of olfactory information transmission is visualized by ODORTRAP and modulated by diet-induced obesity

**DOI:** 10.64898/2026.09.28.755029

**Authors:** Saptarsi Mitra, Franklin A. Pacheco, Debra Ann Fadool

## Abstract

1.

Enhanced excitability of output neurons in the olfactory bulb (OB) has been linked to regulation of energy homeostasis. The mechanism by which the olfactory system can drive metabolic state while processing odor information remains elusive yet must depend upon integration and relay to the rest of the brain. Herein, we screened brain-wide neuronal ensembles that preferentially engaged during odor stimulation (referred to as ODORTRAP) utilizing FOSTRAP activity mapping. Mice were exposed to the odor isopropyl tiglate (IPT), a preferred ligand for the olfactory receptor 160, previously shown to be vulnerable to diet-induced obesity. Odor activation resulted in elevated labelling in the olfactory bulb (ODORTRAP) (*p* < 0.001), as well as in higher olfactory cortices including anterior olfactory nucleus, piriform cortex, and cortical amygdala (*p* < 0.05). Interestingly, IPT differentially recruited hypothalamic feeding areas, namely the arcuate (ARC), dorsomedial (DMH) and lateral (LH) hypothalamus (*p* < 0.001), while the ventromedial hypothalamic nucleus and the suprachiasmatic nucleus exhibited no change in ODORTRAP activity (*p* > 0.05). In mice that were maintained on moderately high-fat (MHF) diets, we discovered that obesity re-models the activation pattern across the brain. Obese mice exhibited loss of ODORTRAP activity in the ARC (p < 0.0001), whereas in the DMH and LH increased baseline activity that obscured any odor-linked neural activity. While IPT activated the nucleus accumbens and medial septum in control mice (*p* < 0.01), obese mice demonstrated no changes in ODORTRAP activity (*p* > 0.05). Our study identifies key brain regions linking chemosensation and metabolic state, specifically demonstrating alteration in activity in hypothalamic domains upon obesogenic diet.

**Keypoints:**

- Investigators used an unique transgenic mouse line that permits Inducible immediate-early gene activation, linking odor stimulation with neural activity (called ODORTRAP) to map circuits beyond the olfactory bulb and olfactory cortices to those of brain regions involved in feeding and energy homeostasis.
- Olfactory input engages hypothalamic feeding-related nuclei – namely the arcuate, the dorsomedial, and the lateral hypothalamus; suggesting a selective integration of olfactory information.
- An obesogenic diet disrupts odor-evoked hypothalamic activation while altering the mesolimbic reward pathway.

## 2. Introduction

Obesity and associated co-morbidities not only impact cardiovascular and metabolic health (de Moura E Dias et al., 2021; Powell-Wiley et al., 2021; Karra et al., 2022; Tutor et al., 2023), they have deleterious effects on brain function. Changes in neuronal activity, specifically in the hypothalamus (Lin and Huang, 1999; Xin et al., 2000; Noronha et al., 2019), structural changes in dendritic spine density (Cope et al., 2018), thickening of extracellular matrix proteins (Beddows et al., 2024), and general reduction in learning, memory, and cognitive flexibility (Davidson et al., 2014; Nguyen et al., 2014; Rukadikar et al., 2023), are a few of the many reported losses attributed to obesity and decreased brain health. Chemosensory systems are particularly sensitive to changes caused by an obesogenic environment (Fadool et al., 2011; Palouzier-Paulignan et al., 2012; Fadool and Kolling, 2020; Harnischfeger and Dando, 2021, 2026). Specifically in the olfactory system, moderately-high fat (MHF) diet consumption has been shown to cause loss of olfactory sensory neurons and their associated axonal projections (Thiebaud et al., 2014; Chelette et al., 2022), changes in microbiome species and inflammatory pathways (Chelette et al., 2022; Loeven et al., 2025, 2026); reduction in cFos immediate-early gene activity in the juxtaglomerular neurons of the olfactory bulb (OB) (Fardone et al., 2019), and dynamic alterations in the transcriptomic profile of olfactory bulb neurons (Kim et al., 2025). These changes culminate to diminished olfactory acuity of the animal; ultimately potentially affecting pre-ingestive as well as feeding behaviours.

Odor-guided sensory perception engages synchronized activity of multiple cell types in the olfactory system as well as the action of circulating metabolites and hormones that reflect the internal energy state of the animal (Dumont et al., 1998; Caillol et al., 2003; Getchell et al., 2006; Tong et al., 2011; Palouzier-Paulignan et al., 2012; Thiebaud et al., 2016; Nogi et al., 2020). Previous reports correlated energy state dependency and olfactory sensitivity, where fasting enhances, and satiety reduces, odor detection thresholds (Aimé et al., 2007, 2014; Wu et al., 2020). There are meal dependent and circadian fluctuations in circulating metabolites such as insulin, leptin, ghrelin, glucagon-like peptide 1 (GLP1), and glucose (citations) that are known to be neuromodulators of olfactory bulb (OB) neurons (Fadool et al., 2000, 2004, 2011; Tucker et al., 2008, 2013; Marks et al., 2009; Thiebaud et al., 2016, 2019; Bell and Fadool, 2017; Huang et al., 2017, 2021; Kolling et al., 2025). The array of metabolite-, neuropeptide– and neurotransmitter-receptor expression in the OB, further confirms how intricately the OB neurons are regulated by these circulating factors (Brünig et al., 1999; Tucker and Fadool, 2002; Caillol et al., 2003; Stanic et al., 2008; Palouzier-Paulignan et al., 2012; Thiebaud et al., 2016; Kovach et al., 2016; Julliard et al., 2017; McIntyre et al., 2017; Bagnoli and FitzGerald, 2019; Fadool and Kolling, 2020; Kolling et al., 2025). This favourably positions the OB as a key homeostatic center beyond the hypothalamus (Leibowitz and Wortley, 2004; Cansell et al., 2012; Tran et al., 2022) to regulate energy balance and food intake in animals. Mitral and tufted cells (M/TCs), the principal output neurons of the OB, form the key node where odor information from olfactory sensory neurons are integrated and relayed to segregated subregions of the olfactory cortex (Igarashi et al., 2012; Nagayama et al., 2014; Kolling et al., 2022). Further refinement of M/TCs output activity is caused by local interneurons in the granule cell layer (GCL; Isaacson & Strowbridge, 1998; Lagier et al., 2007; Thiebaud et al., 2019) as well as centrally secreted neuropeptides and hormones like cholecystokinin, orexin, GLP-1, insulin and glucose (Hardy et al., 2005; Fadool et al., 2011; Economo et al., 2016; Thiebaud et al., 2019; Aghvami et al., 2022; Starr et al., 2025). Upregulation of the excitability of M/TCs using Crispr genome editing or pharmacological block of a major voltage-dependent potassium channel in these neurons, rendered mice resistance to diet-induced obesity by enhancing fuel utilization (reduction of respiratory exchange ratio), improvement of glucose clearance, and alteration of food intake and meal size (Schwartz et al., 2021; Kolling et al., 2022). Thus, alternation of M/TCs excitability acts in a dual capacity to modulate odor information while regulating state of metabolic health.

If enhanced excitability of M/TCs drives metabolic health by changing fuel utilization and oxidation of fats, and by reducing caloric intake and meal size, what connections from the OB to higher cortical regions might drive these physiological changes? It is well known that M/TCs project to overlapping areas in the olfactory cortex such as the anterior and posterior piriform cortex (aPC, pPC), olfactory tubercle (OT), anterior olfactory nucleus (AON), cortical amygdala (CoA), and the lateral entorhinal cortex (LEC) (Scott et al., 1980; Price et al., 1991; Igarashi et al., 2012). The lateral hypothalamus (LH) is a key node that receives direct axonal inputs from the majority of these olfactory cortical subnuclei (Price et al., 1991; Murata et al., 2019; Nogi et al., 2020). Although direct connections between the M/TCs and the hypothalamus are limited, odor information is relayed via intermediary cortical structures like the PC and OT (Price et al., 1991; Bekkers and Suzuki, 2013; Zhang et al., 2017). In addition, polysynaptic tracing studies have revealed connections between the OB and the dorsomedial hypothalamic nucleus (Jovanovic et al., 2023) and the ventromedial hypothalamic nucleus (Liu et al., 2025). The functional link between the OB and the hypothalamus is further informed by studies demonstrating dynamic changes in neural activity in key feeding-related neuronal groups like the agouti-related peptide (AgRP) and proopiomelanocortin (POMC) neurons in the arcuate nucleus (Chen et al., 2015; Su et al., 2017). Conversely, activation of AgRP neurons has been shown to exclusively enhance food odor attraction but not any other olfactory cues (Horio and Liberles, 2021). Brain-wide activity mapping employing conventional c-Fos immunohistochemistry as well as positron emission tomography (PET)-based activity mapping have identified key neural groups like the medial septum VGLUT2 neurons (Bulk et al., 2025), ventral subiculum neurons (Liu et al., 2025), and LH corticotropin releasing hormone (CRH) neurons (Li et al., 2023), which respond to food odors.

Herein, we utilized the approach of Fos-TRAP (Targeted recombination of active populations; (Guenthner et al., 2013; DeNardo et al., 2019)) in conjunction with odor-evoked activity mapping to identify brain regions that respond to odors when mice are maintained on a control, healthy diet vs. that of an obesogenic diet. The TRAP strategy enables permanent genetic labelling of neurons during a narrow activation window (in our case, odor-activation), also providing the scope of future optogenetic or chemogenetic intervention (Guenthner et al., 2013; DeNardo et al., 2019). Despite major progresses in the field of olfactory coding specially at the level of the olfactory epithelium and OB (Junek et al., 2010; Zhou and Belluscio, 2012; Parabucki et al., 2019; Barnum and Hong, 2022), the downstream brain regions activated post odor stimulation, particularly those that are state dependent such as during exposure to obesogenic conditions, are undetermined. Mapping odor-induced TRAP ensembles and comparing across healthy vs. obesogenic diet conditions is the first step in identifying mechanism of how olfactory excitability is linked to whole-body metabolic regulation.

## 3. Materials and Methods

### Solutions and Regents

All general salts and reagents were purchased from Thermo Fisher Scientific, Waltham MA or Sigma Chemical, Atlanta GA unless otherwise noted. Phosphate buffer (PB; 0.1M; pH 7.4) was prepared using equimolar of disodium hydrogen phosphate (Na HPO_4_; Macron Chemicals, Radnor, PA; Cat# 77106) and monosodium dihydrogen phosphate (NaH PO; Thermo Fisher Scientific, Waltham, MA; Cat# S369-500) dissolved in milliQ water. Phosphate-buffered saline (PBS) was prepared by adding 0.1M NaCl (Thermo Fisher Scientific, Cat# 447300010) to PB. Four percent paraformaldehyde (PFA) solution was freshly prepared (Thermo Fisher Scientific, Cat# 416785000) in PB under gentle heat, and adding sodium hydroxide until the solution turned clear. PFA was used fresh or up to 3 days after preparation. Permeabilization Solution (PM) consisted of 0.5% Triton X-100 solution (Thermo Fisher Scientific, Cat# BP151-100) dissolved in PBS with gentle stirring at room temperature (rt). Blocking Solution (BS) was prepared by dissolving 3% normal goat serum (NGS; Jackson ImmunoResearch, West Grove, Pennsylvania; Cat# 005-000-121; RRID: AB_2336990), 0.3% Triton X-100 (Thermo Fisher Scientific, Cat# BP151-100), 0.05% sodium azide (Sigma-Aldrich, St. Louis, MO; Cat# S2002), and 0.2% Kodak Photoflo 200 (Kodak Professional, Rochester, NY; Cat# 1464510) as prepared in PBS. Tris buffer (0.05M) was prepared by dissolving 7.6g Trizma® Pre-set crystals (Sigma-Aldrich, St. Louis, Missouri, Cat# T7693) in 1L millQ water and was used for mounting sections. Isopropyl tiglate (Sigma-Aldrich, Cat# AABH99A42E9D) was dissolved in mineral oil (Sigma-Aldrich, Cat# 330779) to yield a final concentration of 2% and was used for the odor stimulation experiments. Cryoprotection was performed using 10% and 30% sucrose (VWR BDH Chemicals, Radnor, PA; Cat# BDH0308) dissolved in PBS. 4-hydroxytamoxifen (4-OHT; Sigma-Aldrich, Cat#H6278) stock solution was prepared at a concentration of 20 mg/ml dissolved in 100% ethanol. The stock solution was orbitally rotated at 250 RPM at 37°C for 1 h, after which it was aliquoted and stored at –20°C for up to 3 months. On the day of the experiment, frozen aliquots were thawed and orbitally rotated at 250 RPM at 37°C for 30 min. A 1:4 mixture of castor oil (Sigma-Aldrich, Cat# 259853) and sunflower seed oil (Sigma-Aldrich, Cat# S5007) was mixed 50:50 into the 4OHT stock solution, after which the original ethanol solvent was evaporated by vacuum centrifugation to yield a final drug concentration of 20 mg/ml. The final dose (50 mg/kg) was administered intraperitoneally in a volume ranging from 70 to 125 μl depending upon mouse body weight.

### 2.2 Animals

All vertebrate animal procedures were approved by the Florida State University (FSU) Institutional Animal Care and Use Committee (IACUC; Protocol IPROTO202300000001). As such, experiments were done in accordance with the guidelines set by the National Institutes of Health (NIH), the American Veterinary Medicine Association (AVMA), and the ARRIVE guidelines (Percie du Sert et al., 2020). FosTRAP2 mice (B6;129-Fos2A-CreER; JAX stock #030323; The Jackson Laboratory, Bar Harbor, ME; RRID: IMSR_JAX:030323) were crossed with Ai14 tdTomato reporter mice (B6.Cg-Gt(ROSA)26Sor tm14(CAG-tdTomato)Hze /J; JAX stock #007914; RRID: IMSR_JAX:030323) to generate FosTRAP2 x Ai14 (FOSTRAP) mice. Given that the study was designed to examine the variable of fatty diet, we restricted our experiments to males because previous studies have demonstrated that female mice are resistant to diet-induced obesity (Thiebaud et al., 2014; Chelette et al., 2022; Kolling et al., 2022). Animals were housed on a 12 h light/dark cycle (lights on 07:00– 19:00) with food and water available *ad libitum.* The FOSTRAP progeny were weaned at 3 weeks and then randomly assigned to dietary/treatment group, which was initiated within 1 week. Mice were maintained for 6 months on either a control fed (CF) diet consisting of Laboratory Rodent Diet 5001 (LabDiet, St Louis, MO, USA; 28.05% kcal protein, 59.81% kcal carbohydrate and 13.5% kcal fat) or a moderately high-fat condensed milk diet (MHF; Cat# D12266B, Research Diets, Inc., New Brunswick, NJ; 16.8% kcal protein, 51.4% kcal carbohydrate and 31.8% kcal fat). Following dietary treatment, mice were behaviorally tested by stimulating with either mineral oil (control) or odor; see schematic (**Figure 1)**. A total of 23 male mice (body weight ranging from 25-50g) were used in our study.

**Figure 1.**
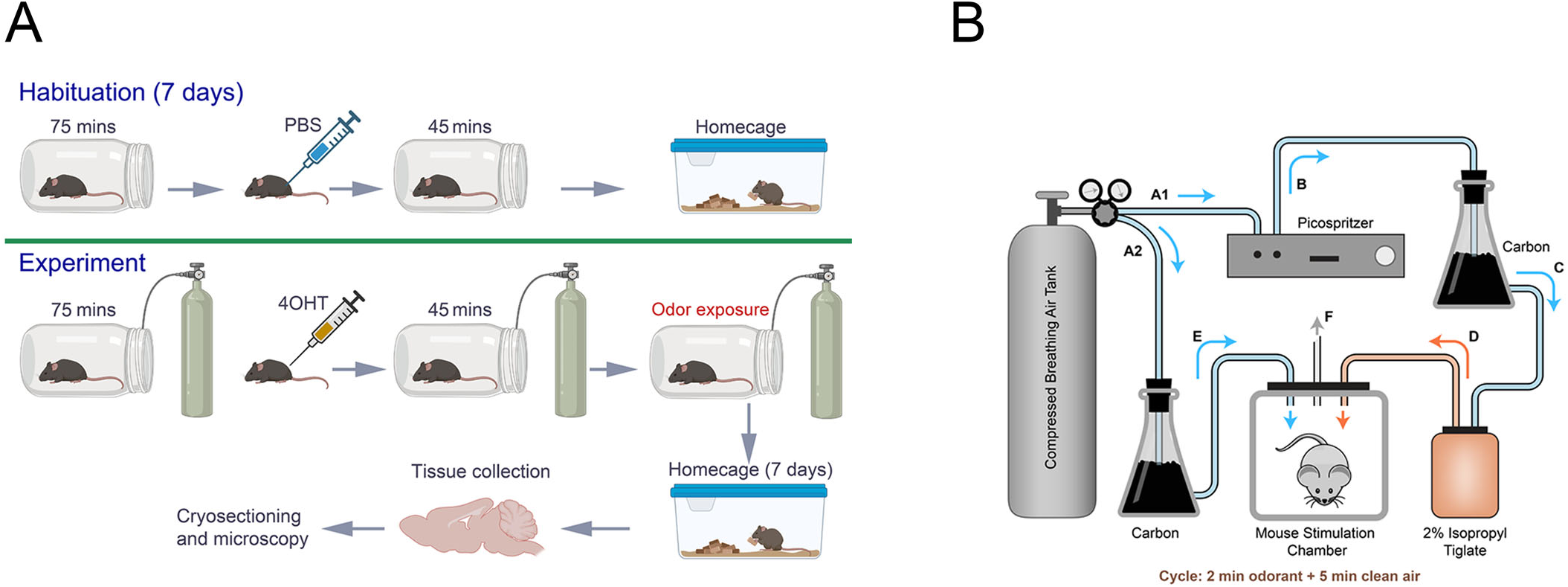
Schematic representation of the experimental procedure. (A) Mice are habituated to the experimental chamber (bell jar) once daily for 7 days prior to the experimental treatment in order to minimize handling and stress from drug administration. After an interval of 75 minutes, mice receive an intraperitoneal (i.p.) injection of PBS followed by retention in the apparatus for 45 minutes prior to be returned to the home cage. On the experimental day, mice alternatively receive an i.p. of 4-hydroxytamoxifen (4OHT) followed by 45 minutes of either odorant or mineral oil (B). Following stimulation, mice are returned to their home cage to allow full tdTomato expression to mark c-Fos gene expression as a proxy for neuronal activity. (B) Details of the odor vs. mineral oil stimulation in a custom-5generated olfactometer. Within the experimental chamber, mice are presented 2-minute cycles of odor/mineral oil followed by 5 minutes of clean air in order to prevent adaption to the stimulus. A total of 4 cycles are delivered over the course of a 28-minute stimulation window. Air is split (A1/A2) from the compressed air cylinder and driven into a (A1) picospritzer to force 20 psi of air into (B) a carbon vessel and enter the (C) odorant or mineral bottle to allow the headspace to enter the (D) experimental chamber to the mouse. After the 2-minute interval applied by the picospritzer, delivery would pause for 5-minutes and only (A2) clean air would be delivered after similarly being scrubbed by (E) carbon.

### 2.3 Habituation, Odor Stimulation, and Tissue Preparation

To minimize handling stress and familiarize the animals to an odor-stimulation chamber, mice were transferred to an experimental room wherein they were habituated to a holding chamber (1L Mason Jars) for 2 h. This was performed daily for 1 week, during which mice received an intraperitoneal (i.p.) injection of sterile PBS approximately half-way into the habituation interval.

On the day of the experiment, animals were stimulated with a patterned frequency of odor using a custom-fabricated olfactometer controlled by a picospritzer (Model II and III, Parker Hanifen Corporation, Hollis, NH; RRID: SCR_018152) that established an air flow at a pressure of 20 psi as described previously (Fardone et al., 2019). Briefly, mice were individually placed in the holding chamber perfused with carbon-filtered clean air for a period of 75 minutes. Following pre-stimulation, the mouse received either an i.p. injection of 4-OHT (drug) or oil solvent (control), and was placed back in the holding chamber for an additional 45 minutes. During this stimulation period, mice were exposed to oscillating patterns of carbon-filtered air (“clean air”) and carbon-filtered air containing mineral oil (control) or 2% isopropyl tiglate (odor). Odor stimulation consisted of four cycles of 2-min odor/mineral oil exposure followed by 5-min clean air. Following stimulation, animals entered a post-stimulation period where they remained in the chamber for an additional 1 h to maximize immediate-early gene (ieg) expression without introduction of additional stimuli. Following the combined stimulation and ieg expression period, mice were returned to their home cage for a period of 7 days to permit sufficient transgene reporter (tdTomato) synthesis.

Brain tissue collection and processing were performed as previously (Mitra et al., 2022). Briefly, mice were anesthetized with ketamine (100 mg/ml, Dechra Pharmaceuticals; Northwich, UK; NDC: 17033-101-10) and xylazine (100 mg/ml, Covetrus; Portland, ME; NDC:11695-4024-1), and transcardially perfused with ice-cold PBS followed by 4% PFA. Brains were post-fixed overnight at 4°C, cryoprotected sequentially in 10% and 30% sucrose, and allowed to completely submerge in solution. The brains were snap frozen in powdered dry ice, mounted in Tissue Tek optimal cutting temperature compound (Sakura Finetek, Tokyo, Japan; Cat# NC1862249) and stored at –80°C until cryosectioned. Serial coronal sections (25 μm thickness) were collected in four alternate sets in a six-well plate. Sections were either processed for immunofluorescence or stored in antifreeze solution (Bioenno Lifesciences, Santa Ana, CA; Cat#006799-250) at –20°C until further processing.

### 2.4 Verification of tdTomato Labeling Specificity

To verify any unwanted background that might be caused by leaky tdTomato expression, we examined conditions for which mice lacked cre-lox capacity. Herein, FOSTRAP mice (without 4-OHT injection) or Ai14 tdTomato reporter mice (lacking inducible fos) were treated with oil solvent (sunflower seed oil + castor oil mixture) or 4-OHT and subsequently exposed to either mineral oil (control) or isopropyl tiglate (odor) as described above. The following combinations of controls were thereby established and are referenced as following –

a) OIL OIL FOS = FOSTRAP mice receiving oil injection and exposed to mineral oil (control).
b) OIL ODOR FOS = FOSTRAP mice receiving oil injection and exposed to isopropyl tiglate (odor).
c) OIL OIL Tomato = Ai1 tdTomato reporter mice receiving oil injection and exposed to mineral oil (control).
d) Drug OIL Tomato = Ai1 tdTomato reporter mice receiving 4-OHT injection and exposed to mineral oil (control)
e) Drug ODOR Tomato = tdTomato reporter mice receiving 4-OHT injection and exposed to isopropyl tiglate (odor) Additionally, we examined FosTRAP2 mice (lacking the Ai tdTomato reporter) to verify that 4-OHT injection is not capable of labelling cells in the absence of the reporter. Namely –
f) Drug OIL FOS = FosTRAP2 mice receiving 4-OHT injection and exposed to mineral oil (control).
g) Drug ODOR FOS = FosTRAP2 mice receiving 4-OHT injection and exposed to isopropyl tiglate (odor).

Brain samples from these animals were similarly processed as mentioned previously and tdTomato labelling was assessed across different brain regions.

### 2.5 Immunofluorescence

To confirm the specificity of odor-induced TRAP (c-Fos gene synthesis) activation, a few coronal hypothalamic sections from odor-exposed FOSTRAP animals were processed for c-Fos antibody immunolabeling. Sections were first rinsed in PBS, immersed in PB for 20 min, and then incubated in BS for 30 min (all at rt). Sections were incubated overnight at 4°C in rabbit monclonal c-Fos antibody (Cell Signalling Technology, Danvers, MA; clone 9F6, Cat #2250; RRID:AB_2247211; 1:1000) diluted in BS. Sections were washed in PBS (3X, 5 minutes/wash), and incubated for 1h at rt with Alexa Fluor 488–conjugated goat anti-rabbit secondary antibody (Thermo Fisher Scientific, Cat #A-11008; RRID:AB_143165; 1:500) diluted in BS. Sections were rinsed in PBS (3X, 5 minutes/wash) and then mounted on VWR® Superfrost slides (Avantor, Cat# 48311-601) in 0.05M TRIS buffer and allowed to dry at rt for 1h followed by coverslipping with Vectashield mounting media containing 4′,6-diamidino-2-phenylindole (DAPI) nuclear stain (Vector Laboratories, Newark, CA; Cat# H1200-10).

### 2.6 Microscopy and Image Analysis

A Keyence BZ-X710 epifluorescence microscope (Keyence Corporation, Osaka, Japan, RRID: SCR_017202) equipped with low power objective lens (4x magnification, 0.1 numerical aperture, and 15.1 mm working distance) was initially used to acquire low-magnification images of the prepared coronal brain sections. Exposure times and gain were maintained consistently across experimental groups. To determine co-expression of tdTomato and c-Fos labelling, photomicrographs were acquired on a Yokogawa Series Nikon Spinning Disk Confocal Microscope (Nikon Instruments, Inc., Melville NY; Model CSUW1, RRID:SCR_021741) using a water-immersion 20x objective lens (0.1 numerical aperture and 15.1 mm working distance). Images were processed using NIS-Elements (Nikon Instruments, Inc.; RRID: SCR_014329) and were compiled using Photoshop CS4 (Adobe, San Jose, CA; RRID:SCR_014199). For quantification of neurons in each brain region, coronal sections (25 μm) were cut throughout the rostrocaudal extent of the brain using a Leica CM2050S cryostat (Leica Biosystems, Wetzlar, Germany, RRID: SCR_016844). Brain regions of interest were identified according to the Allen Mouse Brain Atlas (Lein et al., 2007)). tdTomato-positive neurons were manually counted across different brain regions by the investigator who was blind to the experimental condition. Numerical abundance was tabulated from both hemispheres of the brain using a minimum of 5 mice per experimental treatment group. For each mouse, three to four sections were analysed across each brain area; the data were pooled across sections for the given mouse and brain region of both hemispheres, and the mean ± standard error of the mean (SEM) were determined.

### 2.8 Antibody Characterization

The c-Fos antibody used in this study (rabbit monoclonal C-FOS (9F6, #2250, 1:1,000, Cell Signaling Technology) has been extensively validated. The antiserum was generated using a synthetic peptide corresponding to residues near the amino terminus of human c-Fos protein.

The antiserum detects endogenous levels of c-Fos protein and does not cross-react with other Fos proteins, including FosB, FRA1 and FRA2. This antiserum has been used in a variety of preparations including cultures, frozen sections, and biochemistry experiments. In cortical and hippocampal mixed primary cultures derived from mouse, for example, c-Fos immunoreactivity was observed following bicuculine treatment, but not with vehicle control (Hu et al., 2023). Shang et al. (2023) validated the antiserum by Western blot/SDS PAGE mouse hippocampal lysates at appropriate size of 62 kDa (Shang et al., 2023). In addition, the antibody has been used to reliably label cleared brain samples by iDISCO confirming the accessibility of the antibody in large tissue samples (Branch et al., 2019, bioRxiv). The antibody has also been used to detect neuronal activity across a variety of brain regions including the hypothalamus (Jensen-Cody et al., 2020; Li et al., 2025), the hippocampus (Bjornson et al., 2023) and across cortical areas (Li et al., 2022; Liu et al., 2023). Across these studies, the antibody demonstrated a high degree of nuclear localization and low background labelling. To additionally validate our TRAP-tdTomato labelling, we performed a second round of identical odor stimulation experiments in a TRAP x Ai14tdTomato mouse spaced two weeks apart where co-expression of tdTomato and c-Fos labelling was quantified. This strategy has been previously used by authors to delineate differentially-activated neuronal populations in a brain area during two contrasting/similar stimuli (McDougle et al., 2024; Yang et al., 2025). These studies uniquely identified groups of neurons activated differentially following sugar or fat consumption. The Iba-1 rabbit polyclonal antibody used in our study (019-19741, FUJIFILM Wako Pure Chemical Corporation; RRID:AB_839504) has been previously used and validated in the hypothalamus, basal forebrain, basal ganglia, and insular cortex (Reis et al., 2015; Huang et al., 2020). The antiserum was generated against a synthetic peptide corresponding to the C-terminal sequence of Iba1. We observed increased occurrence of Iba-1 cells in the arcuate nucleus in response to high-fat diet, similar to that of other reports (Valdearcos et al., 2014; Huang et al., 2020). Antibody specificity has been further validated by Western blot analysis from mouse brain tissue lysate that identifies a 17kDa band corresponding to the molecular weight of the protein (Sandhir et al., 2008; Pons et al., 2020)

**TABLE 1.**
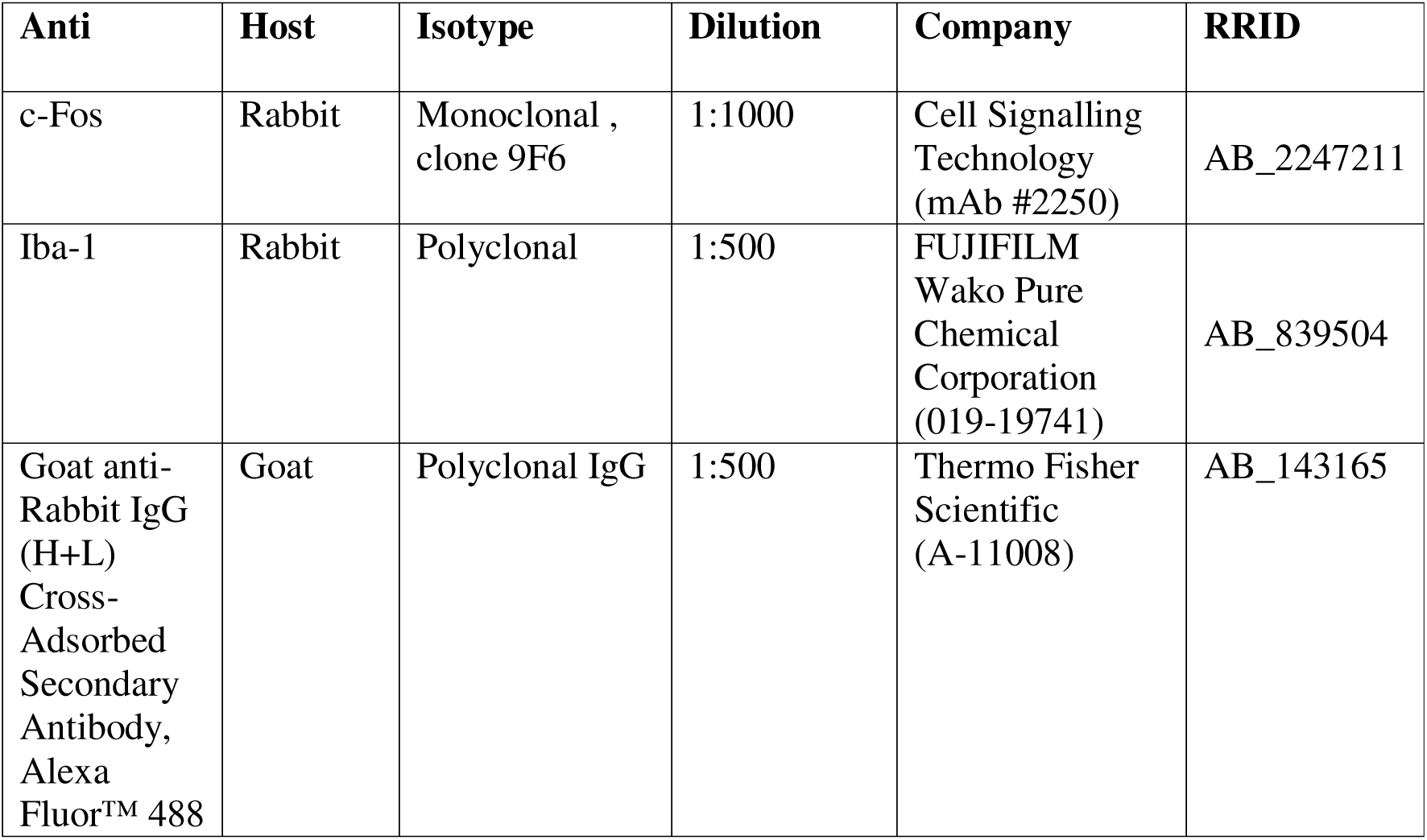
Primary and secondary antibodies used in the study.

### 2.9 Statistical Analyses

All statistical analyses were performed, and graphs were generated using GraphPad Prism 9 (GraphPad Software, San Diego, CA; RRID:SCR_002798). Before performing any statistical comparisons, data were first analysed with the Dixon’s Q test to identify any outliers. Data were then checked for normal distribution and homogeneity of variance using the Fmax test. Analysis of data collected did not identify any outliers. Where collected data violated homogeneity of variance (failed the Fmax test), the selection of applied statistics was changed to the nonparametric equivalent. Data are reported as mean ± SEM and (n) represents number of sampled mice. For pair-wise comparisons across mineral oil vs. odor stimulation treatment groups, a two-tailed Student’s *t* test was performed. For multiple comparisons, a one-way analysis of variance (1-*w* ANOVA) was applied using dietary treatment group as the factor, with a Bonferroni’s *post-hoc* test. Significantly-different means were defined at the 95% confidence interval (α = 0.05). Individual *F* statistic and/or *p* values are reported for each experiment within the corresponding graph and as described in the results section. For 1-*w* ANOVA, not significantly-different means are denoted by the same lower-case letter for clarity of the figures.

## 3. RESULTS

### Activity-dependent neuronal labelling using FOSTRAP approach

To map neural activity across the mouse brain following an odor stimulation experience, we employed the use of Targeted Recombination of Active Populations, using a TRAP2 line containing a tdTomato reporter (FOSTRAP). Because the FOSTRAP line inherently contains an inducible modified cre recombinase (CreERT2) that is dependent upon the presence of the 4OHT ligand, we were able to permanently capture neurons activated during our prescribed odor exposure window. Explicitly during this window, while 4OHT remains active, CreERT2 is able to translocate from the cytoplasm to the nucleus and release the LoxP-Stop-LoxP cassette upstream of the tdTomato reporter (**Figure 2 A1**). Hence, as shown in **Figure 2**, the permanently-labelled neurons can be viewed starting with the olfactory bulb and progressing rostral to caudal in mice that were administered 4OHT and then stimulated with a mineral oil (MO) vehicle control (**Figure 2 A2-A4**) versus that of Isopropyl tiglate (IPT) odor (**Figure 2 B1-B4**). If we repeated the same experiment but substituted the administration of 4OHT with that of sunflower + castor oil (**Figure 2 C1, D1**), then weak, non-specific tdTomato labelling was observed (arrows) in both the OB and brain regions that were not modified by odor stimulation (**Figure 2 C2-C4** vs. **Figure 2 D2-D4**). If we substituted tdTomato mouse lines for that of FOSTRAP lines and administered either sunflower/castor oil or 4OHT (**Figure 2 E1, F1, G1**), no reporter expression was observed under MO– or IPT–stimulated conditions in the absence of the inducible cre recombinase enzyme (**Figure 2 E-G**).

**Figure 2.**
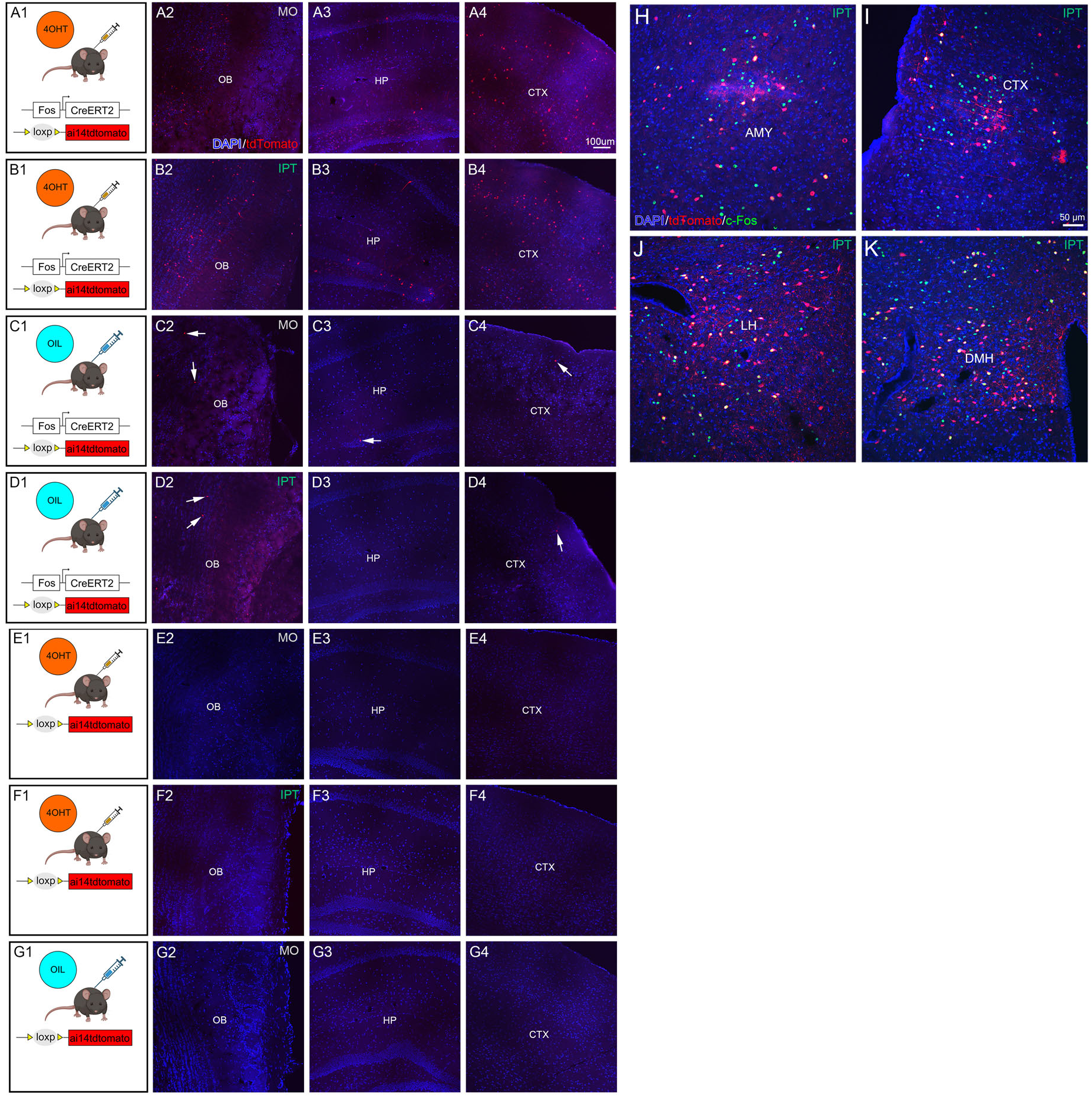
Odor-evoked TRAP labelling following 4OHT-induced synthesis of iCreERT2 that enables excision and permanent tdTomato expression. Representative rostral to caudal brain sections acquired from FOSTRAP mice that were presented (A) mineral oil control (MO) vs. (B) Isoproplyl tiglate odorant (IPT) as demonstrated in Figure 1. A number of control paradigms were undertaken to determine any background expression in the mice in terms of the selected tdTomato reporter: FOSTRAP mice receiving vehicle oil mixture (sunflower + castor) rather than 4OHT, prior to (C) mineral oil (MO) or (D) odorant (IPT) stimulation. tdTomato reporter mice (lacking TRAP ability) rather than FOSTRAP mice, receiving 4OHT prior to (E) mineral oil (MO) or (F) odorant (IPT) stimulation. tdTomato reporter mice receiving vehicle oil mixture (sunflower + castor) prior to (G) mineral oil (MO). Representative coronal sections from isopropyl tiglate (IPT) exposed FOSTRAP mice that underwent TRAP labelling followed by another round of IPT stimulation and subsequent histological processing of brain tissue for c-Fos immunofluorescence. Co-expression of tdTomato and c-Fos imunofluorescence was observed in the (H) amygdala (AMY), (I) cortex (CTX), (J) lateral hypothalamus (LH), and (K) dorsomedial hypothalamus (DMH) Scale bars: 1 mm (A-G); 50 µm (H–K). DAPI = 4′,6-diamidino-2-phenylindole; nuclear stain; tdTomato = reporter for FOSTRAP; c-fos = antibody localization of the immediate-early gene fos.

Next, we examined the degree of co-expression between endogenous c-Fos immunoreactivity and that of FOSTRAP/tdTomato labelling by administering 4OHT as previously described prior to odorant stimulation (**Figure 1A**), and then subsequently stimulating with a second round of odor exposure one week later in the absence of the inducible drug. While the FOSTRAP was anticipated to provide a permanent record of the odor stimulation window, and c-fos antibody detection is known to capture a transient snapshot in time, we still observed a major overlap between tdTomato positive neurons and c-Fos positive cells across different brain regions including the amygdala (AMY), cortex (CTX), lateral hypothalamus (LH), and the dorsomedial hypothalamus (DMH) (**Figure 2J-M**). This result is consistent with the fact that the two approaches have different temporal dynamics, however, the degree of overlap provides consistency in recognizing c-fos as a proxy for neural activity.

### Isopropyl tiglate induces glomerulus-specific and layer specific activation in the olfactory bulb

To assess anticipated glomerular activation within the olfactory bulb we utilized Tbx21-GFP mice that specifically demonstrate GFP expression in mitral and tufted cell bodies and processes. Pairing odor exposure with c-Fos immunocytochemistry revealed a strong glomerular activation, presumably activating periglomerular cells surrounding the OR160-specific glomerulus that is the preferred odorant receptor for the IPT ligand (**Figure 3A-C**).

**Figure 3.**
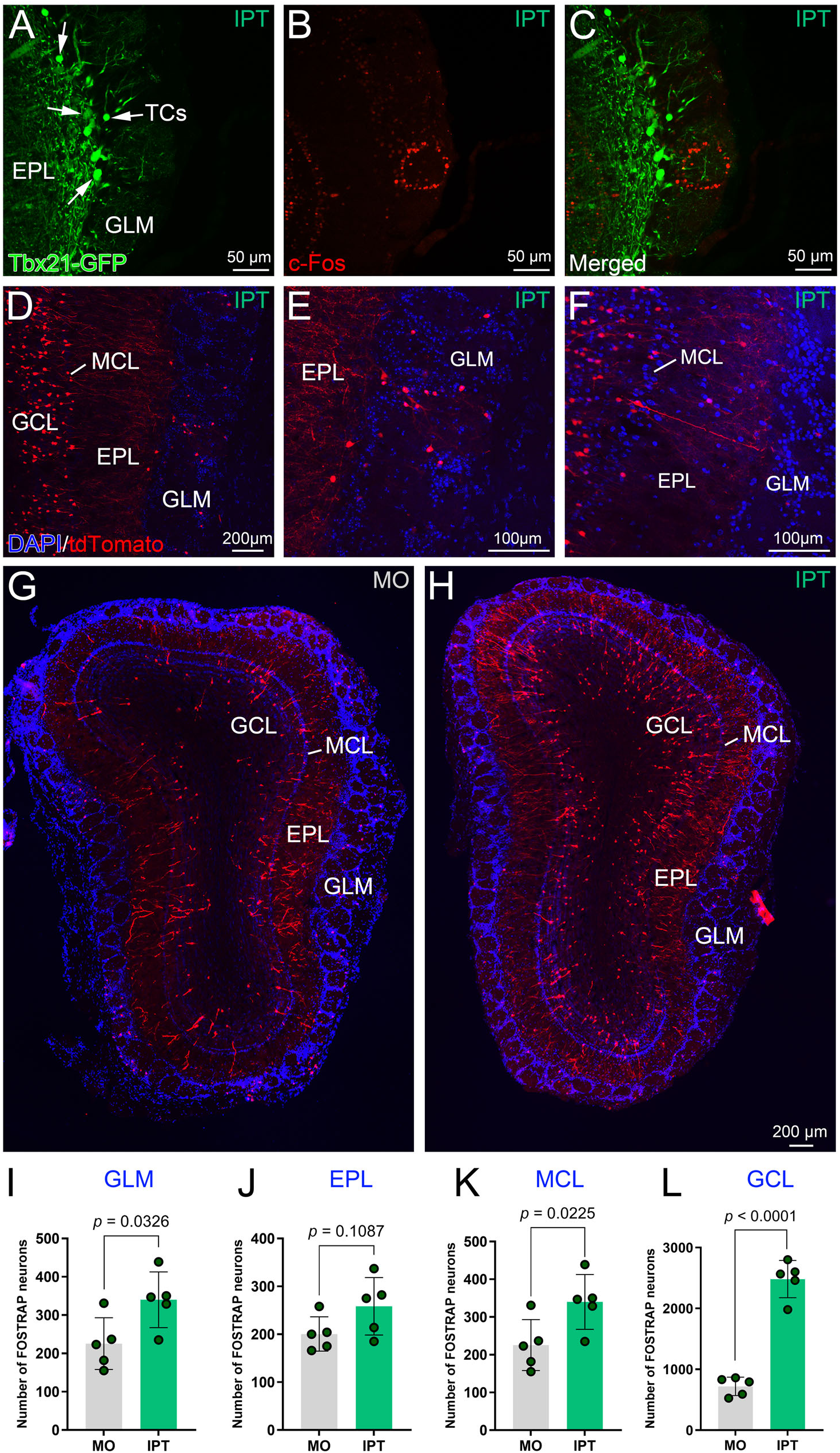
FOSTRAP mice exhibit glomerular-specific activation following isopropyl tiglate (IPT) odor stimulation and have significant elevation of neuronal activity (ODORTRAP) within the glomerular, mitral, and granule cell layers of the olfactory bulb. (A) Representative photomicrograph acquired from aTbx21-GFP mouse demonstrating GFP-expression within the mitral and tufted cells (M/TCs), and (B) c-Fos immunolabeling within periglomerular cells of the activated glomerulus, (C) merged image. Representative photomicrograph acquired from a FOSTRAP mouse demonstrating (D) tdTomato-labelled neurons (red) also within the mitral and tufted cells (M/TCs), and within periglomerular cells of the activated glomerulus, (E) at higher magnification, and (F) showing dendritic projections of the M/TCs. Representative coronal sections acquired from FOSTRAP mice following (G) mineral oil (MO) vs. (H) isopropyl tiglate (IPT) odor presentation. Histogram plots of the number of sampled neurons activated in the (I) glomerular layer (GLM), (J) external plexiform layer (EPL), (K) mitral cell layer (MCL), and (L) granule cell layer (GCL) in IPT-exposed mice compared to that of MO controls. Data are number of mice (n = 5; mean ± SEM; Student’s *t*-test). DAPI = 4′,6-diamidino-2-phenylindole; nuclear stain; tdTomato = reporter for FOSTRAP.

In parallel, the FOSTRAP mice exhibited tdTomato expression within the granule and mitral cell layers that had been only weakly labelled by c-fos immunocytochemistry (**Figure 3D**), however, similar labelling was observed with the tdTomato expression in the periglomerular cells (**Figure 3D-E**) within a specific glomerulus and also mitral cells (**Figure 3F**). Layer specific analyses of the OB revealed significantly increased FOSTRAP labelling in the glomerular layer (GLM; *p* = 0.0326; Student’s *t*-test), the mitral cell layer (MCL; *p* = 0.225; Student’s t-test), and the granule cell layer (GCL; *p* <0.0001; Student’s *t*-test) following IPT exposure compared to that of MO controls, whereas the external plexiform layer (EPL; *p* = 0.1087; Student’s *t*-test) displayed no significant changes in odor-evoked FOSTRAP expression (**Figure 3G–L**; Student’s *t*-test, *p* values reported on the figures for this and subsequent figures).

### Odor-stimulated neuronal activity is wide-spread across the brain

To obtain an overview of the whole brain neural recruitment upon IPT exposure we examined serial coronal sections spanning from the OB through the hindbrain (**Figure 4**). A qualitative approach of estimation was used to analyse fold change in neural activity between the mineral oil– (MO; control) and isopropyl tiglate– (IPT; odor) exposed groups. Analysing olfactory, cortical hypothalamic and limbic brain regions we observed either low, moderate or high degree of changes in activity, while several brain areas showed unaltered levels of neural activity. We understand that odor stimulation recruits variable, yet distinct, neural ensembles across the brain.

**Figure 4.**
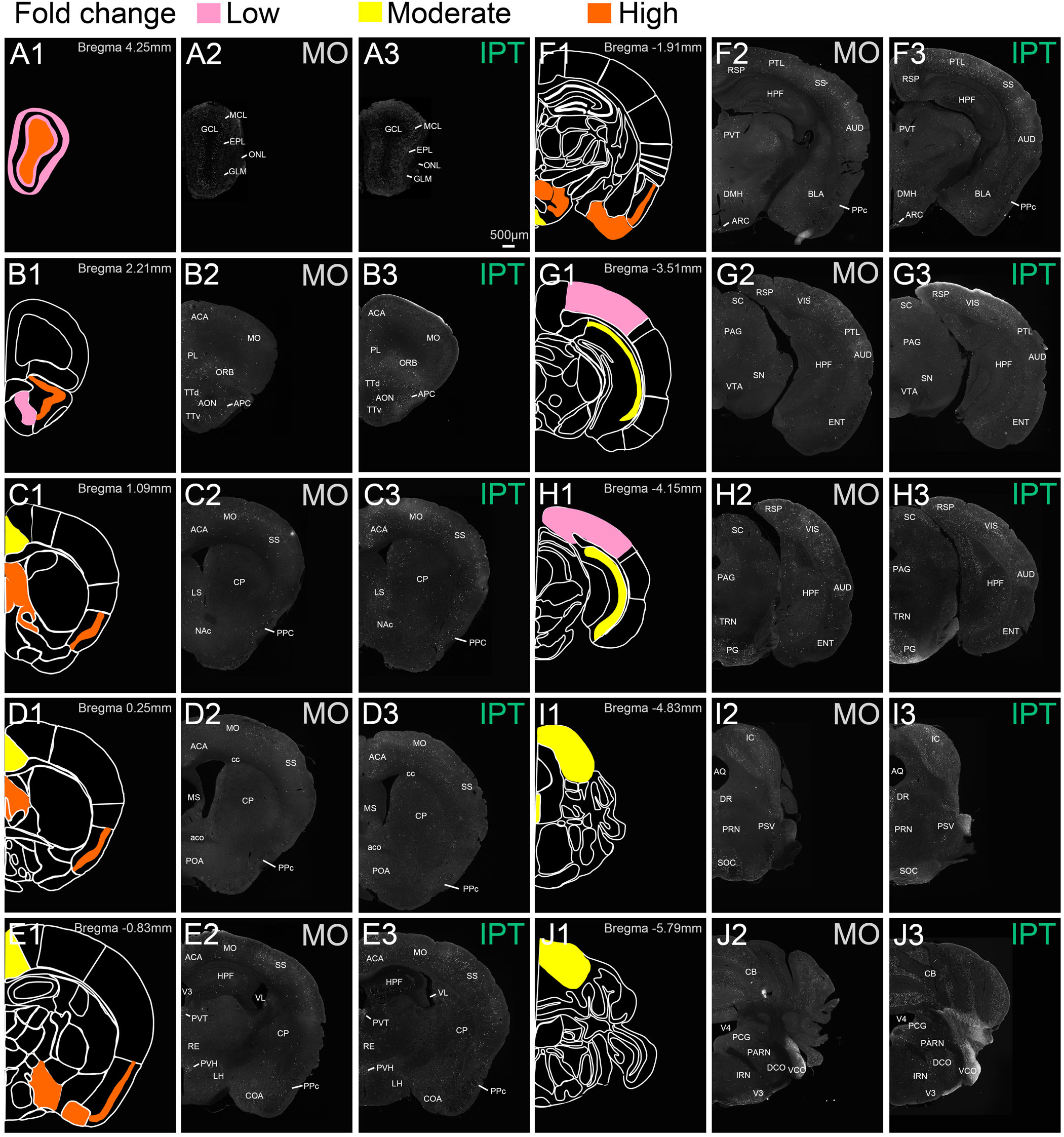
Brain-wide mapping of ODORTRAP activation across the rostrocaudal axis of the brain. Semi-quantitative comparison of ODORTRAP activation using a color-coded indication of fold increase of IPT/MO change in tdTomato expression. Within each row set (example A1-A3), the first panel is the color-coded fold increase of IPT/MO change overlaid across the Allen Brain Map template (A1). The second panel is a representative coronal section following MO presentation (A2) vs. that of the third panel that is following IPT presentation (A3). Low (pink), moderate (yellow), or high (orange) levels of tdTomato expression. Scale bar = 500 µm. *Abbreviations:* ACA, anterior cingulate area; NAc, nucleus accumbens; AON, anterior olfactory nucleus; AQ, cerebral aqueduct; APc, Anterior piriform cortex; ARC, arcuate hypothalamic nucleus; AUD, auditory area; BLA, basolateral amygdalar nucleus; CB, cerebellum; COA, cortical amygdala; CP, caudoputamen; DCO, dorsal cochlear nucleus; DMH, dorsomedial hypothalamic nucleus; DR, dorsal nucleus raphe; ENT, entorhinal area; EPL, external plexiform layer; GCL, granule cell layer; GLM, glomerular layer; HPF, hippocampal formation; IC, inferior colliculus; IRN, intermediate reticular nucleus; LH, lateral hypothalus; LS, lateral septum; MO, Primary motor area; MCL, mitral cell layer; MS, Medial septum; ONL, olfactory nerve layer; ORB, orbital area; PAG, periaqueductal gray; PARN, parvicellular reticular nucleus; PCG, pontine central gray; PG, pontine gray; PPc, posterior piriform cortex; PL, prelimbic area; POA, preoptic area; PRN, pontine reticular nucleus; PSV, principal sensory nucleus of the trigeminal; PTL, posterior parietal association area; PVH, paraventricular hypothalamic nucleus; PVT, paraventricular nucleus of the thalamus; RE, nucleus of reuniens; RSP, retrosplenial area; SC, superior colliculus; SN, substantia nigra; SOC, superior olivary complex; SS, somatosensory area; TRN, tegmental reticular nucleus; TTd, dorsal tenia tecta; TTv, ventral tenia tecta; V3, third ventricle; V4, fourth ventricle; VCO, ventral cochlear nucleus; VIS, visual area; VL’ Lateral ventricle; V4, 4th ventricle, VTA, ventral tegmental area.

### Odor-stimulated neuronal activity recruits downstream olfactory cortical and amygdalar sites

Given the strength of the neuronal activity in the olfactory bulb, we quantified the immediate downstream targets (**Figure 5**). The dorsal and ventral tenia tecta (dTT and vTT) of the primary olfactory cortex have direct connections with the olfactory bulb important for odor-guided behaviors and odor perception, therefore it was surprising that neither showed significant odor-evoked increased activity (**Figure 5A-E**; TTd; *p* = 0.4529; Student’s t-test, and TTv; *p* = 0.2073; Student’s t-test). IPT stimulation, however, significantly increased FOSTRAP labelling in the anterior olfactory nucleus (AON; *p* = 0.0131; Student’s t-test) and the anterior piriform cortex (APC; *p* <0.0001; Student’s t-test; **Figure 5I-J**). Robust FOSTRAP activation was observed in the posterior piriform cortex (PPC; *p* < 0.0001; Student’s t-test), especially within layers II and III containing principal pyramidal cells (**Figure 5F-H, N-O**) – indicating the engagement of the olfactory cortex during IPT exposure. Finally, in these downstream olfactory regions, strong activation was observed in the cortical amygdala (COA; *p* = 0.0003; Student’s *t*-test) (**Figure 5K-N)**

**Figure 5.**
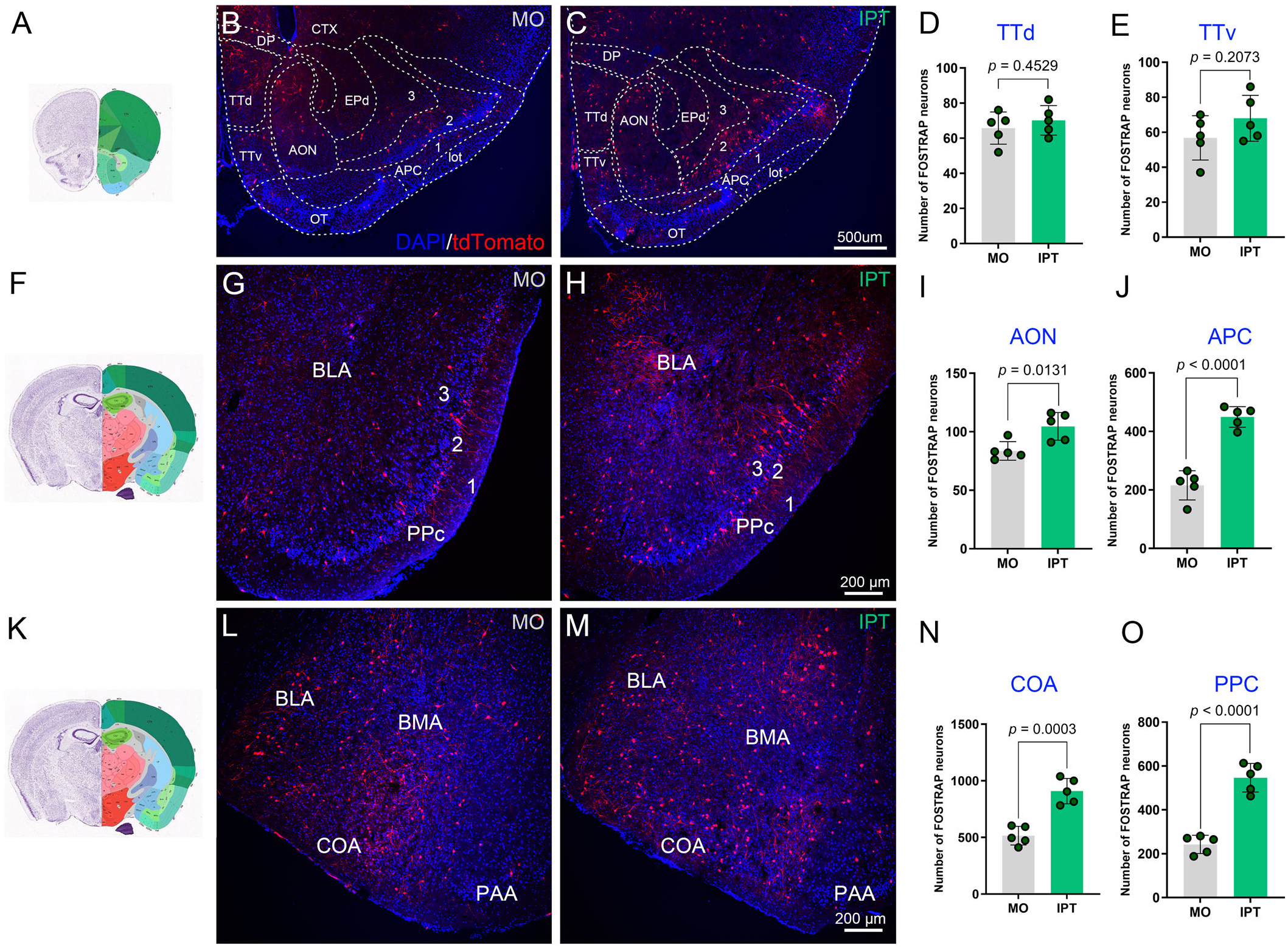
ODORTRAP activity is observed in the accessory olfactory bulb, the posterior piriform cortex (PPC) and the basolateral amygdala (BLA). Representative coronal sections acquired from FOSTRAP mice (Rostral to Caudal; (A, F, K)) following (B, G, L) mineral oil (MO) vs. (C, H, M) isopropyl tiglate (IPT) odor presentation. Histogram plots of the number of sampled neurons activated in IPT-exposed mice compared to that of MO controls located in the (D) dorsal tenia tecta (TTd), (E) ventral tenia tecta (TTv), (I) anterior olfactory nucleus (AON), (J) anterior piriform cortex (APC), (N) cortical amygdala (COA), and (O) posterior piriform cortex (PPC), Layers I, II, III as indicated. Data are number of mice (n = 5; mean ± SEM; Student’s *t*-test). DAPI = 4′,6-diamidino-2-phenylindole; nuclear stain; tdTomato = reporter for FOSTRAP.

### The paraventricular thalamus and preoptic area remain unaffected by odor exposure

IPT exposure did not alter FOSTRAP expression in the paraventricular nucleus of the thalamus (PVT; *p* = 0.8311; Student’s *t*-test) compared to that of MO controls (**Figure 6A– D**). Similarly, FOSTRAP activation in preoptic area subdivisions, namely, anterodorsal preoptic nucleus (ADP; p = 0.8950; Student’s *t*-test), anteroventral periventricular nucleus (AVPV; p = 0.7996; Student’s *t*-test), and anteroventral preoptic nucleus (AVP; p = 0.3315; Student’s *t*-test) were also unaltered by odor presentation compared with that of MO (**Figure 7A–D**). This expression pattern suggests that odor cues alone are insufficient to recruit thalamic or preoptic neural subsets that are associated with either sexual pheromone-based activation (Martinez and Petruslis, 2013) or which are sensory hubs for energy state dependency (Livneh et al., 2017).

**Figure 6.**
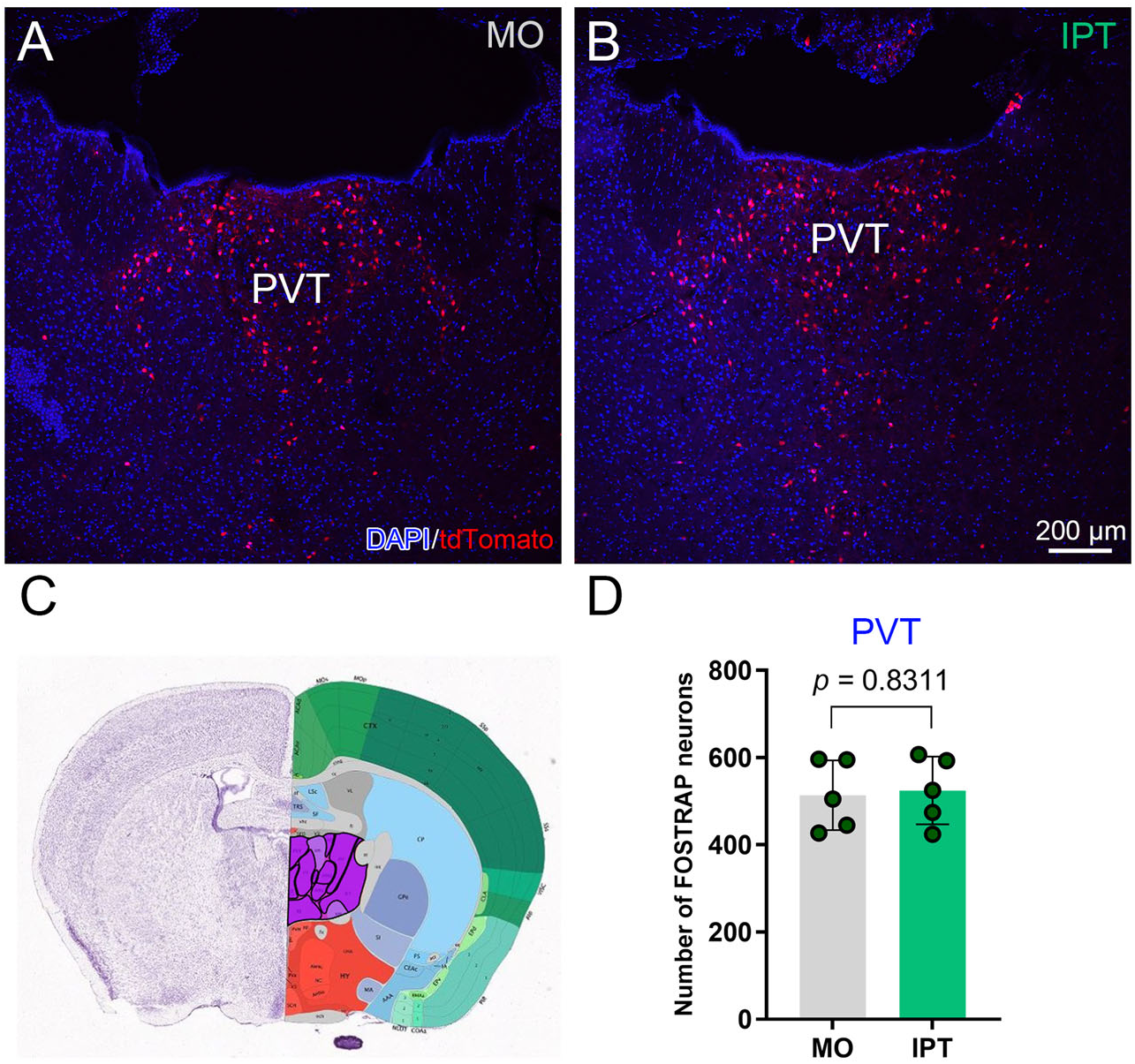
Isopropyl tiglate exposure resulted in unaltered neural activity in the paraventricular thalamic nucleus (PVT). Representative coronal sections acquired from FOSTRAP mice following (A) mineral oil (MO) vs. (B) isopropyl tiglate (IPT) exposure. (C) Atlas plate illustrating the anatomical location of the paraventricular thalamic nucleus (PVT). (D) Histogram plot of the number of sampled neurons in MO vs. IPT groups. Data are number of mice (n = 5; mean ± SEM; Student’s t-test). DAPI = 4′,6-diamidino-2-phenylindole; nuclear stain; tdTomato = reporter for FOSTRAP.

**Figure 7.**
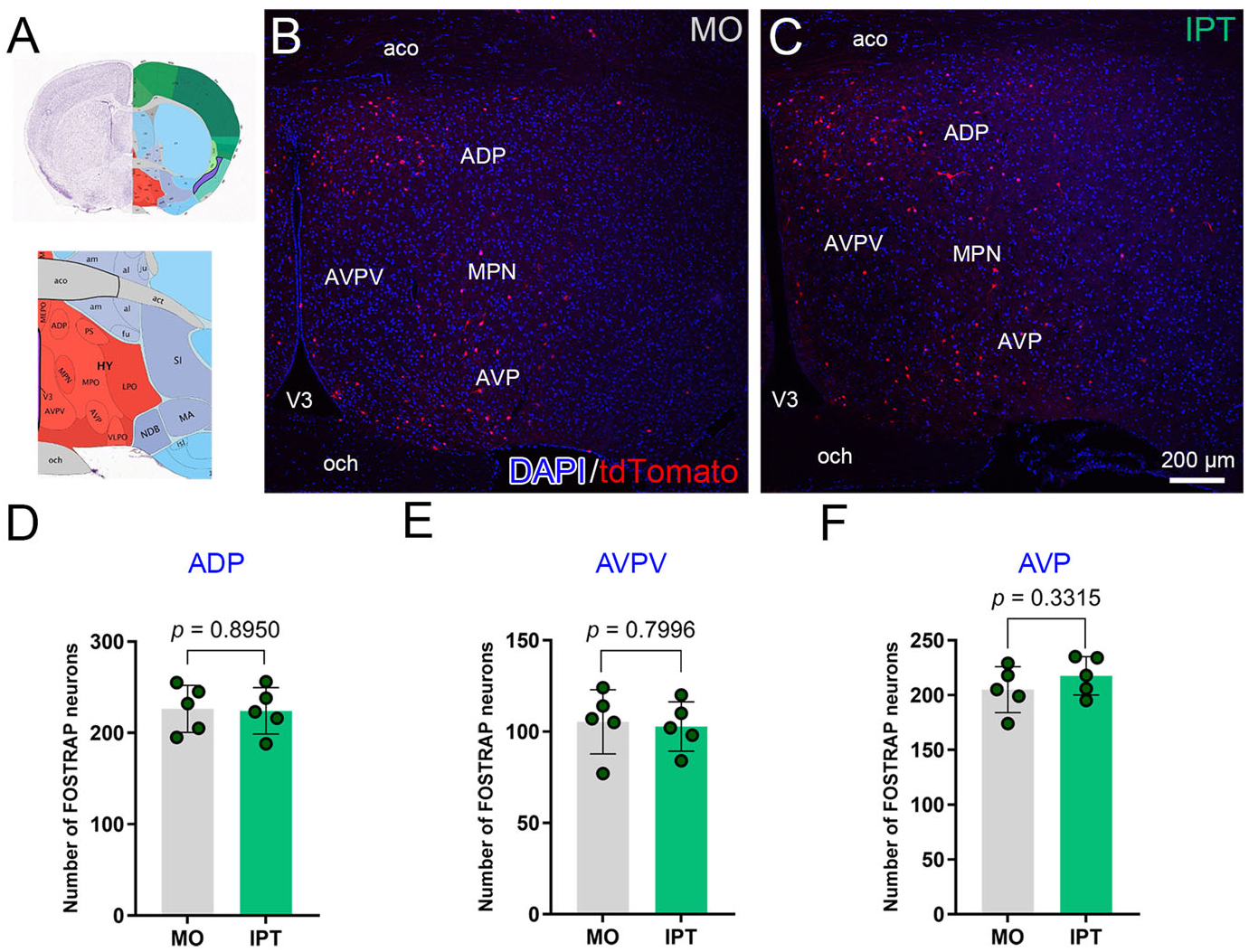
Separate preoptic area subdivisions demonstrate no changes in FOSTRAP activity upon isopropyl tiglate exposure. (A) Low and high magnification atlas plates showing the location of distinct preoptic area subdivisions; anterior dorsal preoptic nucleus (ADP), anteroventral periventricular nucleus (AVPV), medial preoptic nucleus (MPN), and anteroventral preoptic nucleus (AVP). Coronal sections acquired from (B) mineral oil (MO) and (C) isopropyl tiglate (IPT) exposed mice demonstrating no significant changes in tdTomato expression. Histogram plots of the number of sampled neurons activated in the (D) ADP, (E) AVPV and (F) AVP of IPT – exposed mice compared to that of MO controls. Data are number of mice (n = 5; mean ± SEM; Student’s *t*-test). DAPI = 4′,6-diamidino-2-phenylindole; nuclear stain; tdTomato = reporter for FOSTRAP.

### Odor stimulation selectively engages hypothalamic feeding nuclei

We next homed in on the hypothalamic regions given the association of odor molecules with eating and the known link of olfactory activity to metabolic balance. Odor stimulation significantly increased FOSTRAP labelling over that of MO stimulation in feeding-related nuclei, including the arcuate nucleus (ARC; *p* = 0.0019; Student’s *t*-test), the lateral hypothalamus (LH; *p* < 0.0001; Student’s *t*-test), and the dorsomedial hypothalamus (DMH; *p* < 0.0001; Student’s *t*-test) (**Figure 8A-D, M-P, U-X**). FOSTRAP labelling was not significantly increased following odor stimulation in several other hypothalamic regions including the ventromedial hypothalamus (VMH; *p* = 0.2907; Student’s *t*-test), the paraventricular hypothalamus (PVH; *p* = 0.1294; Student’s *t*-test), or the suprachiasmatic nucleus (SCN; *p* = 0.4235; Student’s *t*-test) (**Figure 8E–L, Q–T**). Absence of FOSTRAP expression in the VMH is consistent with a lack of stress-associated neurons being engaged by the IPT stimulation (Nguyen et al., 2024).

**Figure 8.**
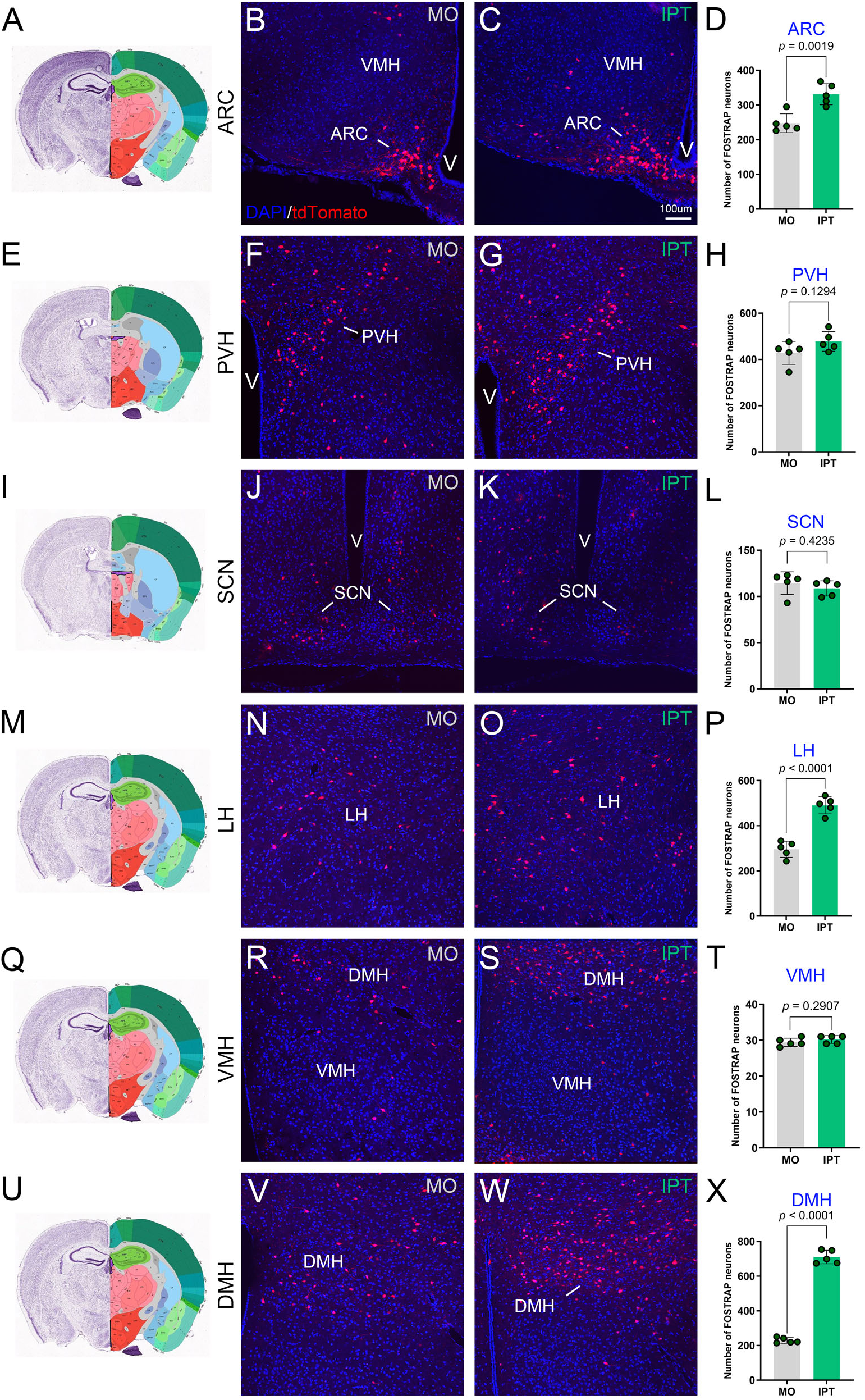
Odor stimulation engages TRAP activity in distinct hypothalamic nuclei. Atlas plates (A, E, I, M, Q, U) depicting the anatomical location of the different hypothalamic nuclei and their corresponding photomicrographs from mineral oil (MO) and isopropyl-tiglate (IPT) exposed groups demonstrating tdTomato expression in the (B-C) arcuate nucleus (ARC), (F-G) paraventricular hypothalamic nucleus (PVH), (J-K) suprachiasmatic nucleus (SCN), (N-O) lateral hypothalamus (LH), (R-S) ventromedial hypothalamic nucleus (VMH) and (V-W) the dorsomedial hypothalamus (DMH). Histogram plots of the number of sampled neurons across these different hypothalamic nuclei of IPT-exposed mice compared to that of MO controls (D, H, L, P, T, X). Data are number of mice (n = 5; mean ± SEM; Student’s *t*-test). DAPI = 4′,6-diamidino-2-phenylindole; nuclear stain; tdTomato = reporter for FOSTRAP.

### Dietary state modifies FOSTRAP expression in the hypothalamus

Because excess consumption of dietary fat causes a loss of olfactory sensory neurons and associated axonal projections (Thiebaud et al., 2014; Chelette et al., 2022), we surmised that diet-induced obesity would cause a functional loss of FOSTRAP odor-evoked activity, particularly that observed for the ARC, LH, and the DMH. We therefore examined FOSTRAP activation in the hypothalamus and compared cohorts of mice that were maintained on control (CF) vs. moderately high-fat diet (MHF) for 6 months post weaning. We refined our analysis to examine the anterior and caudal subdivisions of this brain region separately. Consistent with what we observed for odor-activated FOSTRAP mice across the ARC, both the anterior and caudal ARC subdivisions had significant increases in labelling when mice were maintained on CF diet (**Figure 9A, B, F, I, J, Q, R**; caudal = p < 0.0001, anterior = p < 0.0001; 1-w ANOVA, Tukey’s post-hoc test). When mice were alternatively maintained on the MHF diet, however, there was not only a lack of increased FOSTRAP labelling with odor stimulation over that of mineral oil (**Figure 9C, D, F, K, L, S, T**; caudal = *p* > 0.9999, anterior = *p* > 0.9999, 1-w ANOVA, Tukey’s post-hoc test) but there was a reduced basal labelling in the MO condition in general compared with that of mice maintained on CF diets (**Figure 9A, C, F**; caudal = *p* < 0.0001, anterior = *p* < 0.0001, 1-w ANOVA, Tukey’s post-hoc test). For the IPT-induced activation of the DMH and LH, both the anterior and caudal subdivisions had significantly increased FOSTRAP labelling over that of MO in CF-maintained mice (**Figure 9A, B, H, G, I, J, O, P**; DMH caudal = *p* < 0.0001, DMH anterior = *p* < 0.0001, LH caudal = *p* < 0.0001, LH anterior = *p* < 0.0001, 1-w ANOVA, Tukey’s post-hoc test), however, in the MHF-maintained mice, there was a significant elevation in basal labelling in the MO controls compared to CF-maintained mice, that the odor stimulation was completely obscured (i.e. not significantly different than MO). In fact, the MO-stimulated animals had similar to greater FOSTRAP labelling in the MHF-maintained mice than that of odor-stimulated, CF-maintained mice. These data suggest that diet-induced obesity selectively disrupts odor-driven recruitment of ARC neurons while in the DMH and LH, odor-activated signalling is completely masked by high basal activity.

**Figure 9.**
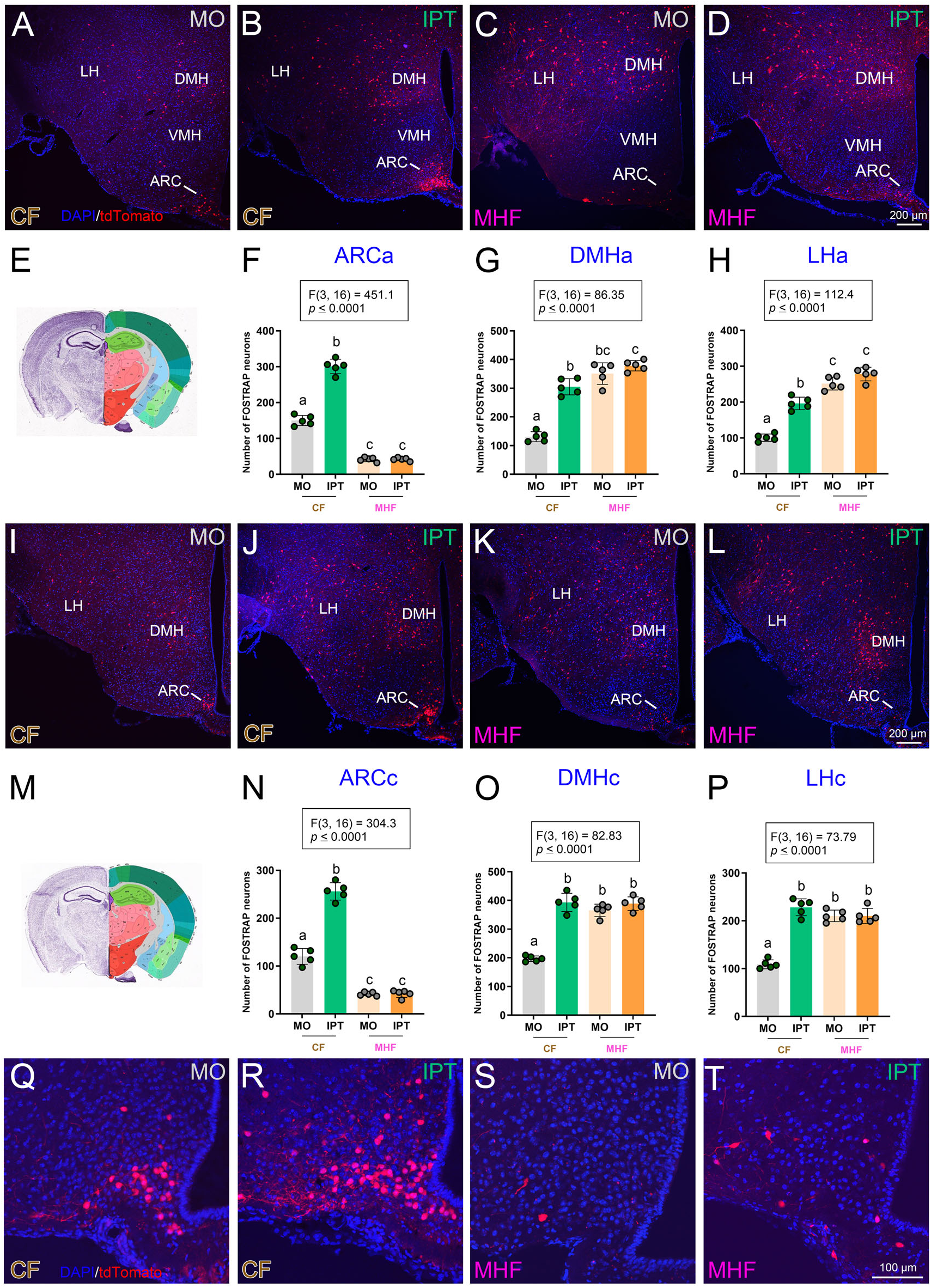
Selective alteration of hypothalamic neural activity upon moderately-high fat diet intervention. Representative photomicrographs coronal sections acquired from FOSTRAP mice maintained on control-fed (CF) diet following (A, I) mineral oil (MO) vs. (B, J) isopropyl tiglate (IPT) odor presentation and from mice maintained on moderately high-fat (MHF) diet following (C, K) mineral oil (MO) vs. (D, L) isopropyl tiglate (IPT) odor presentation. Atlas plates illustrating the anatomical location of the anterior (E) and caudal (M) hypothalamic regions. Histogram plots of the number of sampled neurons activated in the anterior (F) arcuate nucleus (ARCa), (G) dorsomedial hypothalamus (DMHa), and (H) lateral hypothalamus (LHa), and the caudal (N) arcuate nucleus (ARCc), (O) dorsomedial hypothalamus (DMHc), and (P) lateral hypothalamus (LHc). Higher magnification representative photomicrographs of the arcuate nucleus from CF mice following (Q) mineral oil (MO) vs. (R) isopropyl tiglate (IPT) odor presentation and MHF mice following (S) mineral oil (MO) vs. (T) isopropyl tiglate (IPT) odor presentation. Data are number of mice (n = 5; mean ± SEM; 1-*w* ANOVA with Tukey’s *post-hoc* test). Significantly-different means are denoted by different lower-case letters. Scale bars = 200 µm (A–D, I–L); 100 µm (Q–T).

### Alteration of mesolimbic reward pathway in mice maintained on moderately high-fat (MHF) diet

We next examined the ventral tegmental area (VTA) and nucleus accumbens (NAc) that constitute brain areas within the mesolimbic reward pathway (Keifflin et al., 2019).

Regardless of diet, IPT stimulation failed to significantly increase FOSTRAP labelling over that of MO stimulation within the VTA or SNc (**Figure 10A, D, G, J, F, I**; VTA; *p* = 0.8320; SNc; *p* = 0.6350; 1-*w* ANOVA, Tukey’s *post hoc* test). In contrast, IPT stimulation evoked a robust increase in FOSTRAP neurons in the NAc of CF maintained mice (**Figure 10B, E**, L; *p* < 0.0001; 1-*w* ANOVA, Tukey’s *post hoc* test), which was absent in MHF maintained mice, despite comparable baseline FOSTRAP labelling under MO conditions (**Figure 10 H, K, L;** *p* = 0.9217; 1-*w* ANOVA, Tukey’s *post hoc* test). These findings suggest that diet-induced obesity selectively disrupts the sensory-driven activation of the reward pathway. Moreover, this corroborates previous findings showing loss of OSNs during MHF dietary intervention in mice (Thiebaud et al., 2014), which appears to prevent downstream activation of neural sub-circuits.

**Figure 10.**
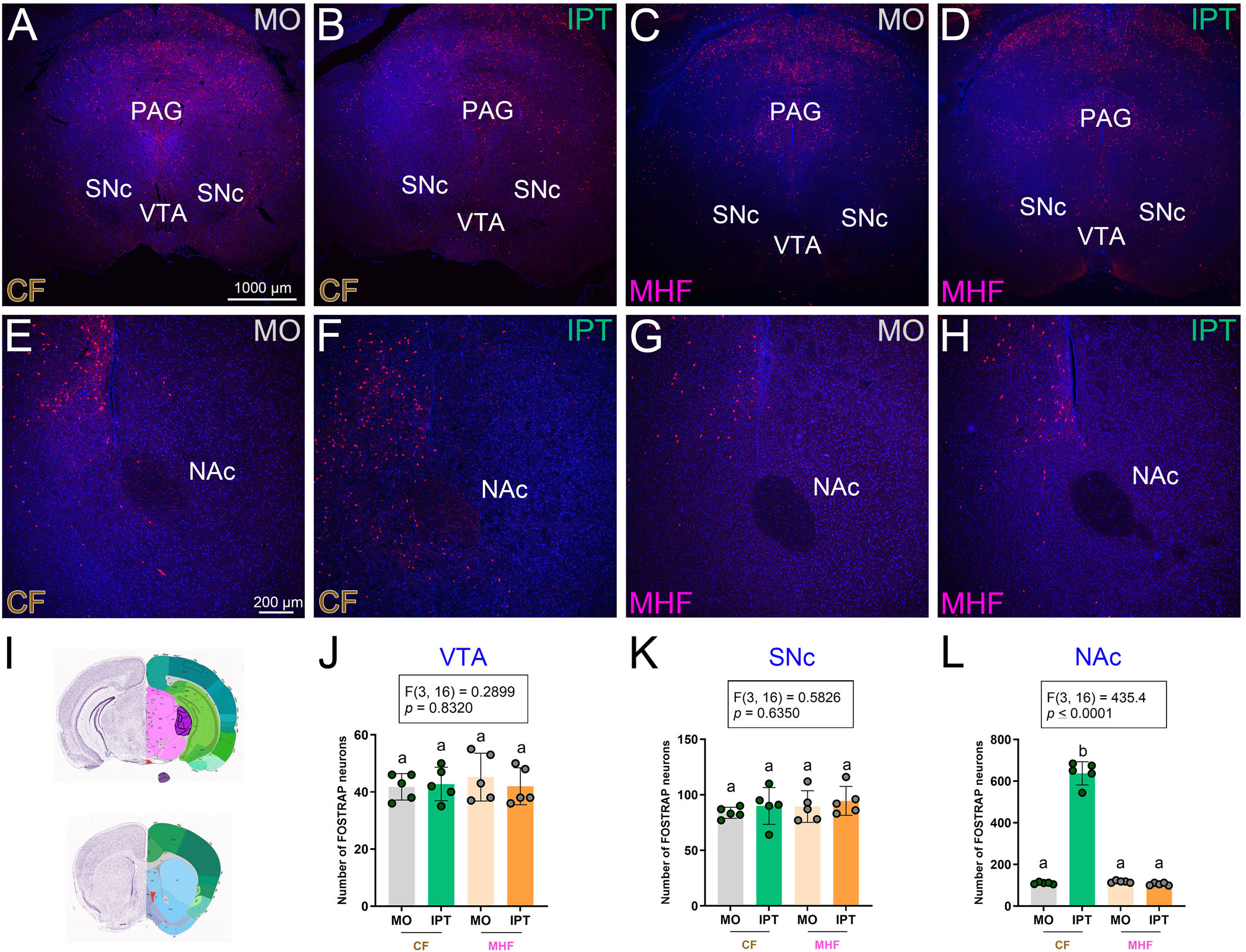
Moderately high-fat diet alters odor-evoked neural activity in the nucleus accumbens but not the midbrain. Representative coronal sections acquired from FOSTRAP mice maintained on control-fed (CF) diet following (A, B) mineral oil (MO) and (D, E) isopropyl tiglate (IPT) odor presentation and from mice maintained on moderately high-fat (MHF) diet following (G, H) mineral oil (MO) vs. (J, K) isopropyl tiglate (IPT) odor presentation. (C) Atlas plates illustrating the anatomical location of the ventral tegmental area (VTA), substantia nigra pars compacta (SNc), and nucleus accumbens (NAc). Histogram plots of the number of sampled neurons activated in the (F) ventral tegmental area (VTA), (I) substantia nigra pars compacta (SNc), and (L) nucleus accumbens (NAc). Data are number of mice (n = 5; mean ± SEM; 1-*w* ANOVA with Tukey’s *post-hoc* test). Significantly-different means are denoted by different lower-case letters. Scale bars = 1000 µm (A, D, G, J); 200 µm (B, E, H, K).

### Odor–evoked septal activation is blunted by diet-induced obesity

Given recent evidence implicating the role of the medial septum (MS) in food odor driven anticipatory activity, we quantified FOSTRAP labelling in this region following IPT stimulation. FOSTRAP labelling was significantly increased following IPT stimulation compared to that of MO controls for mice that were maintained on CF diets (**Figure 11A, B, F**; *p <* 0.0001; 1-*w* ANOVA, Tukey’s post hoc test). In contrast, following diet-induced obesity, IPT stimulation did not increase FOSTRAP labelling over that of controls nor was basal activity elevated to mask any change in neural activation (**Figure 11C, D, F**; *p* = 0.6043; 1-*w* ANOVA, Tukey’s post hoc test). This pattern is supported by previous reports by Bulk et al., 2025 who demonstrated that medial septum neurons are selectively recruited by food-predictive olfactory cues in lean animals to support anticipatory motivational drive, and that this recruitment is markedly affected during high-fat consumption.

**Figure 11.**
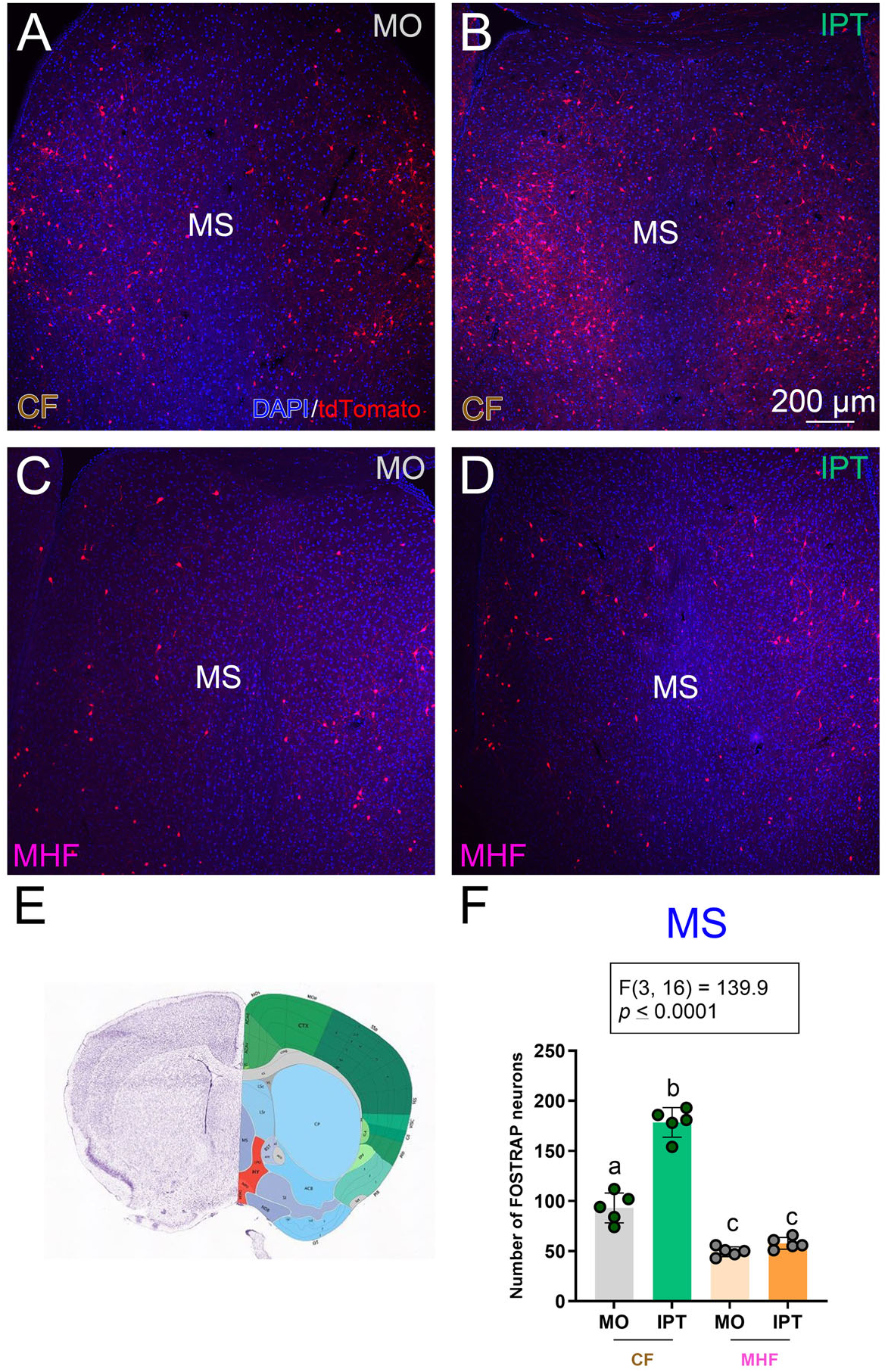
Moderately high-fat diet blunts FOSTRAP activity in the medial septum independent of odor exposure. Representative coronal sections acquired from FOSTRAP mice maintained on control-fed (CF) diet following (A) mineral oil (MO) vs. (B) isopropyl tiglate (IPT) odor presentation and from mice maintained on moderately high-fat (MHF) diet following (C) mineral oil (MO) vs. (D) isopropyl tiglate (IPT) odor presentation. (E) Atlas plate illustrating the anatomical location of the medial septum (MS). Histogram plot of the number of sampled neurons activated in the medial septum (MS) (F). Data are number of mice (n = 5; mean ± SEM; 1-*w* ANOVA with Tukey’s *post-hoc* test). Significantly-different means are denoted by different lower-case letters.

### Increased microglial recruitment in the arcuate nucleus of moderately high-fat diet mice

It is known that diet-induced obesity affects glial morphology attributed to metabolic stress (Douglass et al., 2017). We therefore employed the microglial marker Iba-1 to examine the structure of glia in the ARC because it labels both active and resting states. Compared to that of CF-maintained mice, MHF-maintained animals exhibited an increase in Iba-1 positive glial cells **(Figure 12A, B)**. Moreover, microglia in MHF-maintained animals appeared more numerous and dense throughout the ARC, consistent with previous reports (Huang et al., 2020). We also observed Iba-1 positive microglial processes in close apposition to FOSTRAP-labelled neurons in the ARC in MHF mice (**Figure 12E**), indicating enhanced neuron-microglia interactions following diet-induced obesity. These findings are consistent with previous reports demonstrating that high-fat diet exposure significantly increases both the number and cell body size of Iba-1–positive microglia within the arcuate nucleus (Huang et al., 2020).

**Figure 12.**
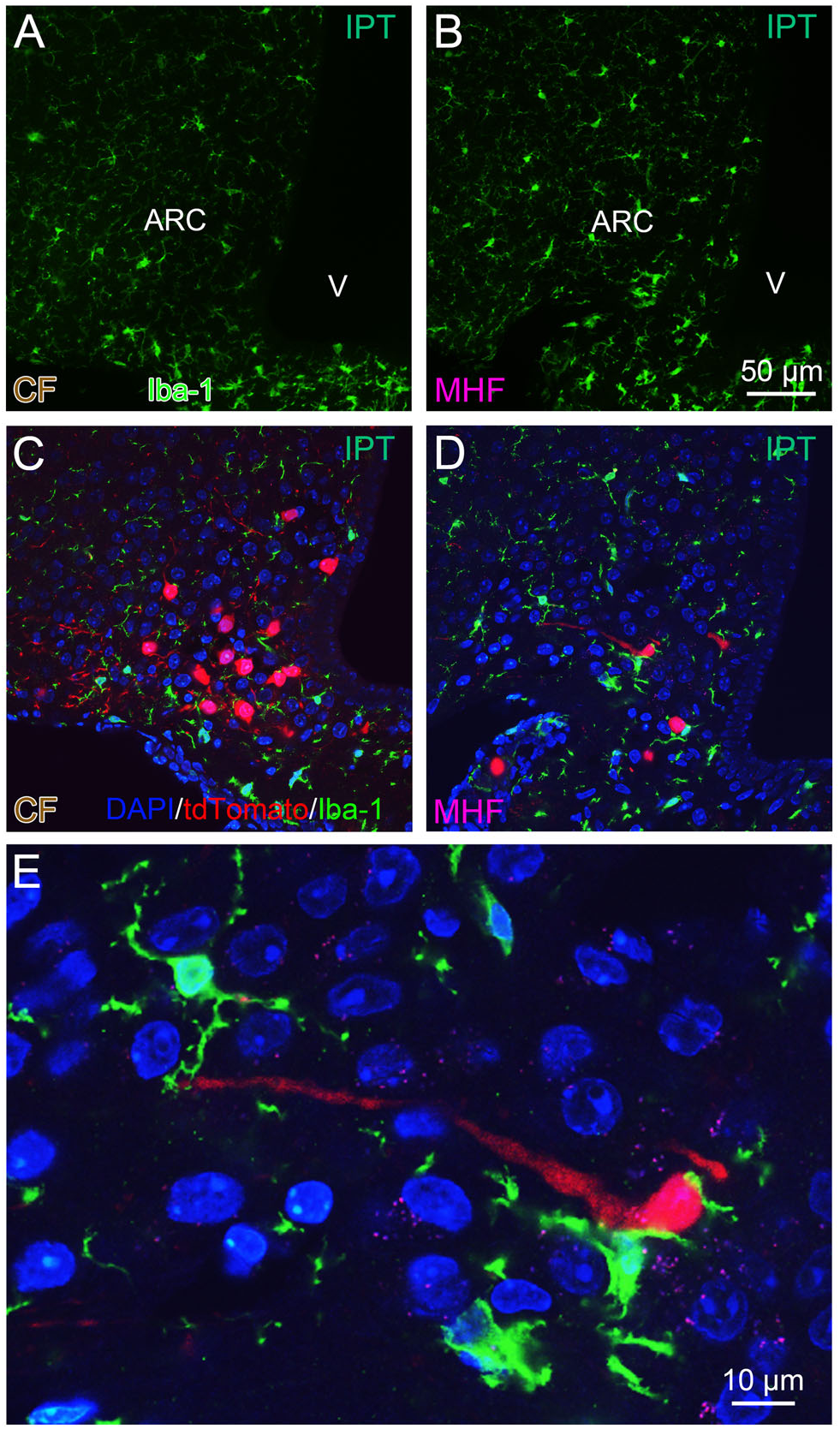
Elevated microglial expression in the arcuate nucleus of moderately high fat exposed animals. Representative photomicrographs demonstrating Iba-immunofluorescence (green) in the arcuate nucleus (ARC) of isopropyl-tiglate (IPT) exposed mice maintained on either (A) control-fed (CF) or (B) moderately high-fat (MHF) diet and their corresponding association with TRAP labelled tdTomato neurons within the ARC in CF (C) or MHF (D, E) exposed animals. Scale bars = 50 µm (A–D); 10 µm (E).

## 4. DISCUSSION

We optimized and employed the FOSTRAP strategy to permanently label neurons that were activated by the preferred ligand of the olfactory receptor 160 (Olfr160), isopropyl tiglate (IPT), in control fed (CF) and moderately high fat (MHF) fed animals. Henceforth we will refer to these permanently-labelled neurons as ODORTRAP. Along with an anticipated activation pattern across the olfactory cortex, odor stimulation distinctly, and interestingly, activated the hypothalamus. Animals that underwent MHF diet intervention, demonstrated selective blunting of neural activity in the medial septum (MS), the nucleus accumbens (NAc) and the arcuate nucleus (ARC). These findings indicate that odor stimulation not only activates brain regions regulating odor coding but simultaneously relays to centers of metabolism and energy homeostasis. Moreover, the extent of odor activation is modulated depending upon the metabolic state of the animal and is dampened with excess energy consumption.

Because this was the first use of FOSTRAP strategies in the olfactory system, we were systematic in handing of the mice to minimize immediate early-gene activation that was irrelevant to the behavioral activation, in our case odor stimulation. As such, we established a controlled habituation paradigm where the animals were handled daily, introduced to glass jars that mimicked the odor chamber (Fardone et al., 2019), and received a daily control intraperitoneal injection for 7 days to habituate the animal to both the injection/handling stress and the novelty of the odor chamber. A comparable habituation paradigm to mitigate stress-induced neuronal responses has been demonstrated in previous reports (Leake et al., 2021; Stoltenborg et al., 2022). We also wanted to verify the specificity of the TRAP-tdTomato labelling by analysing the co-expression of tdTomato and c-Fos immunofluorescence signals in odor-activated brain regions. Thus, a few TRAP2-tdTomato mice underwent odor stimulation with 4OHT, but also received a 2^nd^ round of odor stimulation 2 weeks later followed by processing brain tissue for c-Fos immunofluorescence. Herein, we observed a high degree of co-expression of tdTomato and c-Fos in diverse brain regions such as the amygdala, cortex and hypothalamus. A similar strategy that combines TRAP-tdTomato labelling and c-Fos immunofluorescence has been previously utilized to identify neurons in the preoptic area that responded to thirst (Allen et al., 2017), and differentially activated neurons responding to either sugar or fat in the nucleus tractus solitarius (McDougle et al., 2024) and the hippocampus (Yang et al., 2025). Our comparison of control experiments in both TRAP2 and tdTomato reporter strains further demonstrated that our signal was highly attributed to odor stimulation and not containing high leaky reporter expression that would obscure the true signal. CREer^T2^ strains do demonstrate low levels of 4OHT-independent recombination in neurons resulting in reporter expression in the absence of neuronal activity (Guenthner et al., 2013; DeNardo et al., 2019) similar to what we observed in our study. The low, comparable level of signal across both MO– and IPT-exposed groups could be quantified and subtracted in the analysis as independent of odor-evoked neural activity. The TRAP2 line that we utilized in our study is notably the second generation of CreERT2 lines, which demonstrate improved recombination efficiency (DeNardo et al., 2019) compared to that of the 1^st^ generation TRAP1 lines (Guenthner et al., 2013).

We first quantified ODORTRAP activity within the olfactory system for which the chemosensory stimulus was targeted. We commenced our observations with the olfactory bulb rather than the olfactory epithelium where G-protein-coupled odor transduction is initiated, because fos is not expressed to a detectable extent in the periphery (Loch et al., 2015). There was a focal activation in the GLM of Tbx-21-GFP mice (Kolling et al., 2022, 2025), in addition to weakly-activated glomeruli that were presumably periglomerular neurons (Johnson et al., 1995; Fardone et al., 2019) as estimated by both c-Fos immunolabelling and TRAP-tdTomato labelling. The GLM engagement is potentially dependent on the IPT concentration and odor receptor-specific cascade activation (Vincis et al., 2012). Mitral cells (MCs) are under dual regulation from a spatially-distributed network of glomeruli that selectively exert both excitatory and inhibitory control on mitral cells (Fantana et al., 2008). This supports our findings where we observe the presence of multiple glomerular activations presumably regulating a distinct set of functionally similar MCs.

Mineral oil (MO), due to its low volatility and relatively neutral smell, has been a widely-used diluent for odor experiments in rodents (Moreno et al., 2009; Fardone et al., 2019). Mice can effectively detect and discriminate even between different brands of mineral oil (Gamble and Smith, 2009), thus it was important for us to quantify baseline TRAP activity in parallel mice that were presented simply control mineral oil. The mineral oil would activate an olfactory sensory neuron (OSN)-mediated transduction cascade, which would be propagated to the OB (Nagayama et al., 2014), despite it being the control solvent. In terms of odor stimulation, IPT-mediated activation of neurons in the GLM indicates both OSN recruitment via Olfr160 and also the concentration of the odorant molecule (Rubin and Katz, 1999; Vincis et al., 2012; Zhou and Belluscio, 2012). MC activation underscores that IPT stimulation was sufficient to propagate odor information to downstream brain regions (Nagayama et al., 2014). Additionally, enhanced GC activity that is predominantly inhibitory interneurons in the OB (Nunes and Kuner, 2018; Takahashi et al., 2018), further fine-tunes the M/TC output (Cavarretta et al., 2016), indirectly affecting output sites of the OB. MC activation can further release glutamate onto GC dendrites creating a lateral inhibition loop (Pimentel and Margrie, 2008). Unlike c-Fos immunolabelling, TRAP-tdTomato labelling is not restricted to the nucleus. Interestingly, a dense plexus of GC dendrites and axons can be seen terminating onto M/TCs, hence giving us a clear picture of activated GC subtypes potentially regulating M/TC activity. Importantly, use of IPT as the odorant is supported by prior work identifying it to be a preferred ligand of the Olfr160 (Potter et al., 2001) with-diet-induced changes in c-Fos activity in the Olfr160-expressing glomeruli (Fardone et al., 2019), further underscoring the circuit specificity and metabolic dependence of the activated neurons (Biju et al., 2008; Chelette et al., 2022).

As a consequence of elevated ODORTRAP activity of the principal output neurons of the OB (M/TCs), we next sought to investigate the immediate monosynaptic output targets of the OB. Tracing studies have previously revealed temporally-distinct projection patterns of M/TCs, with each cell type terminating in distinct subfields in the olfactory cortex. While TCs terminate into more anterior divisions of the olfactory cortex, MCs innervate both immediate and distant subfields like the PC and COA (Igarashi et al., 2012; Nagayama et al., 2014). In line with these evidences, IPT stimulation evoked significant increases in ODORTRAP labelling within the AON, which is implicated in integration of MC output response (Lei et al., 2006) and also storage of olfactory memories (Aqrabawi and Kim, 2020). Previous odor stimulation experiments have revealed differential contribution of the pars externa and pars principalis subdivisions of the AON in facilitating cue dependent responses (Kay et al., 2011). AON activation therefore depicts novelty of the IPT stimulation and subsequent integration and storage of this neural information originating from the main OB. In contrast, both the dorsal and ventral tenia tecta (D/VTT) subdivisions did not show significant alteration in ODORTRAP activity compared to that of the matched MO controls. The VTT has been implicated in learned, goal-directed behaviour (Shiotani et al., 2020), thus it may not be anticipated to engage these neurons after a single delivery of IPT with no prior associations to a reward with the odor. In view of the stimulus-specific activity of the DTT (Cousens, 2020), exposure to IPT alone might not have been sufficient to trigger significant changes in neural activity in this area. IPT exposure produced robust ODORTRAP labelling across both anterior (APC) and posterior (PPC) piriform cortex (PC) subdivisions. The APC, a primary projection field of M/TCs, encodes odorant molecular features and fine sensory discriminations between odor combinations (Litaudon et al., 2003; Gottfried et al., 2006; Howard et al., 2009; Stettler and Axel, 2009; Bolding et al., 2020). In contrast, the PPC receives comparatively less dense innervation from the OB but is heavily innervated by the amygdala and the entorhinal cortex, in turn governing aspects of odor memory and contextual aspects of odor (Chapuis et al., 2013; East et al., 2021; Majak et al., 2004; Pedroncini et al., 2024). Finally, the cortical amygdala (CoA) is also heavily innervated by the MCs and is implicated in innate odor-evoked behaviors, particularly odor valence, instinctive olfactory behaviors (Root et al., 2014), and spatial odor identity, in contrast to the distributive representations observed in PC (Miyamichi et al., 2011; Sosulski et al., 2011). Together, this effectively positions the CoA as a critical interface linking olfactory sensory input with downstream hypothalamic and motivational circuits.

Stemming out from the anticipated higher olfactory cortical regions, we next wanted to understand the patterns of ODORTRAP activity found beyond these projections of general olfactory information. MO– and IPT-exposed animals demonstrated comparable ODORTRAP activity across the different subdivisions of the preoptic area (POA). While POA is known to regulate sleep (Mondino et al., 2021; Rothhaas and Chung, 2021; Maurer et al., 2024), thermoregulation (Boulant, 2000; Zhao and Riters, 2023), and reproductive behaviour (d’Anglemont de Tassigny et al., 2010; Glanowska and Moenter, 2015), evidence underscores the involvement of POA, particularly the MPN and AVPV subdivisions, in processing socio-reproductive pheromone cues (Wood and Newman, 1995; Coolen et al., 1998; Swann et al., 2001; Fewell and Meredith, 2002). Since IPT closely resembles the smell of cherry and is a non-reproductive cue, we speculate segregated pathways distinctly regulating food odor– and reproductive-cue processing.

Within the paraventricular nucleus of the thalamus (PVT) the ODORTRAP signal was also highly equivalent between solvent vs. odor stimulation. Both MO and IPT represent novel olfactory stimuli to the animals because they never had previous experience to either. We speculate that each could potentially trigger the arousal/novelty-based neuronal network in the PVT. This is consistent with the established role of the PVT as a salience and an arousal hub integrating hypothalamic and brainstem inputs with limbic motivational circuits (Ren et al., 2018; Barson et al., 2020). The PVT has also been shown to dynamically track behavioral arousal transitions and environmental responsiveness (Setzer et al., 2022), while molecular profiling studies further highlight its involvement in regulating motivational and behavioral processes (Gao et al., 2023)

ODORTRAP activity within the hypothalamus differentially labelled structural nuclei governing eating behaviors, energy balance, and response to circadian-driven endocrine factors. IPT exposure resulted in significant activation of the feeding-related nuclei: the arcuate nucleus (ARC), the lateral hypothalamus (LH), and the dorsomedial hypothalamus (DMH), while homeostatic and circadian regions: the paraventricular nucleus of the hypothalamus (PVH), the ventromedial hypothalamus (VMH), and the suprachiasmatic nucleus (SCN) showed no differential changes in activity across control– and IPT-exposed animals.

The IPT exposure that demonstrated a significant increase in ODORTRAP activity in the ARC was quite focal yet may be complex to interpret. Chen et al. (2015) have used selective expression of calcium indicators within the AgRP and POMC neurons to demonstrate that the sensory detection of food odor is sufficient to engage, and then reset, circuits within the ARC (Chen et al., 2015). Optogenetic activation of ARC AgRP neurons has been conversely shown to increase attraction to food but not to pheromone odors (Horio and Liberles, 2021). Although AgRP neuronal activity reduces as soon as food odor is presented (Betley et al., 2015; Chen et al., 2015), no such changes occur when false food smell is presented (Betley et al., 2015). IPT is a synthetic molecule (sweet, minty odor) and does not have the complex properties of food, such as the aroma from chow, which may alter the inhibition normally experienced with food odor and associated ARC neuronal activity.

We haven’t investigated the cell-type neurochemical properties yet, hence our odor-evoked FOSTRAPPED neurons might be a heterogeneous population of ARC neurons comprising both anorexigenic and orexigenic populations. Additionally, it is important to note that the FOSTRAP method permanently captures neuronal activity over a period of time, hence the ARC FOSTRAP neurons give us a spatial signal but not necessarily temporal information following the IPT presentation. Hence, it is possible that IPT odor exposure in the absence of immediate food access, particularly in sated animals, potentially failed to suppress AGRP activity, and instead caused activation of these neurons.

We judge that the heightened ODORTRAP labelling we observed in the LH likely represents odor-evoked engagement potentially for the initiation of feeding or motivational drive. The LH receives multi-synaptic olfactory input from the olfactory bulb and olfactory peduncle (Murata et al., 2019; Jovanovic et al., 2023). Conversely, we and others have reported reciprocal LH-orexin and LH-MCH projections that may support bidirectional modulation with the OB (Qi et al., 2023, 2026). Functionally, LH neuronal subtypes, including corticotropin-releasing hormone (CRH) and orexin populations, are activated by food cues and are known to drive feeding motivation and consummatory behaviors (Sakurai et al., 1998; Stuber and Wise, 2016).

Finally, within the hypothalamus, ODORTRAP mapping revealed a broad and robust increase in neuronal recruitment within the DMH. The DMH functions as an integrative hub regulating feeding, energy expenditure, torpor, circadian output, and arousal through interactions with broader hypothalamic and limbic networks (Chou et al., 2003; Singru et al., 2005; Ambler et al., 2022). Importantly, recent circuit-mapping studies demonstrate polysynaptic connectivity linking OB to DMH, providing an anatomical pathway through which olfactory sensory signals can access the DMH (Jovanovic et al., 2023).

Other areas within the hypothalamus and HPA axis, showed no change in ODORTRAP activity. PVH is a major neuroendocrine and autonomic integrative hub regulating energy expenditure, stress responses, and satiety through diverse neuronal populations, including CRH, TRH, and oxytocin and vasopressin neurons (Herman and Tasker, 2016; Sutton et al., 2016; Qin et al., 2018; Iremonger and Power, 2025). Given the fact that PVH neurons are mainly under the regulation of internal energy state and hormones, and that GLP1R expressing neurons in the PVH have been shown to be rapidly activated by food odor (Li et al., 2019), it was surprising that this area of the inner and upper hypothalamus was not engaged. Likewise, we did not observe differential ODORTRAP labelling in the SCN across MO– and IPT-exposed animals. We were careful to conduct all odor-exposure experiments within a few hours of the dark cycle because it is well-known that the OB has a dedicated circadian clock with olfactory sensitivity peaking during the active period in mice (Granados-Fuentes et al., 2006), however despite the SCN serving as both the primary circadian regulator (Coomans et al., 2015; Hastings et al., 2018) and its projections to feeding centers in the hypothalamus (Zhai et al., 2022; la Fleur et al., 2024), this region was not odor-activated. ODORTRAP labelling in the VMH also lacked significant odor-evoked activity. The VMH is strongly implicated in processing predatory cues via vomeronasal pathways (Carvalho et al., 2020). SF1+ neurons mediate defensive and fear responses to aversive olfactory stimuli (Kunwar et al., 2015). The absence of differential TRAP activation therefore suggests inherent property of the IPT odor, which does not engage the vomeronasal fear pathway.

Within the three feeding-related hypothalamic nuclei – ARC, LH, and DMH – that each demonstrated robust, ODORTRAP activity, diet-induced obesity muddled the activation of these brain circuits. Within the ARC, we observed desensitization in both control– and IPT-stimulated obese mice compared to those that were maintained on control diet, and within the DMH and LH, the baseline FOSTRAP activity was so elevated that no odor-evoked change in activity could be discerned. Wang et al. (1999) and Xin et al (2000) have previously demonstrated that high-fat diet elevates baseline c-Fos activity in the DMH and LH, thus lack of odor-evoked activation in our study could represent baseline neuronal activity associated with the MHF diet, rather than lack of odor-evoked recruitment of these neurons. Nonetheless neural activity information could not be encoded with the changed state of obesity. These data may alternatively suggest that the effect of chronic MHF feeding may extend beyond a simple reduction in hypothalamic odor responsiveness but rather reflect an altered sensory processing by selectively disrupting odor-evoked recruitment of ARC neurons. This possibility aligns with previous literature, showing that a chronic high-fat diet may disrupt synaptic processes within the ARC, including a reduction in excitatory postsynaptic current amplitude in ARC neurons (Wang and Li., 2022), suggesting that dietary exposure can remodel the synaptic properties of neurons involved in energy homeostasis.

These changes in excitatory synaptic transmission could alter the strength of inputs received by ARC neurons and may contribute to the lack of odor-evoked activity observed following MHF feeding. OB influence over ARC activity must be indirect, mainly due to the polysynaptic route of communication mediated by intermediate sensory and limbic areas (Kuang et al., 2026). The desensitization of ARC neurons in MHF animals in both control– and IPT-exposed animals could be attributed to inflammatory signalling and leptin/insulin resistance within ARC neurons (Ullah et al., 2021), obesity-associated extracellular matrix remodelling around ARC neurons restricting peripheral accessibility and hormonal signalling (Beddows et al., 2024), or long-lasting desensitization of AgRP neurons in the ARC during obesity (Beutler et al., 2020). Previous studies from our laboratory have revealed collateral loss of neurons in the olfactory epithelium and their associated olfactory projections into glomeruli (Thiebaud et al., 2014), as well as changes in glomerular excitability (Fardone et al., 2019) and mitral cell firing frequency (Fadool et al., 2011) in response to obesity, which would further reduce upstream ensemble excitability innervating the ARC.

An interesting observation regarding the hypothalamic response following MHF feeding was that diet-induced obesity did not produce a uniform change in neuronal activity across feeding-related nuclei. Instead, the difference between basal– and odor-evoked activity appeared to be altered in a region-dependent manner. For instance, in the ARC, DIO was associated with a reduction in baseline labelling and a lack of increased ODORTRAP labelling following odor stimulation compared to MO, whereas odor-evoked signalling is completely masked by an elevated basal FOSTRAP labelling in the DMH and LH. Therefore, these data suggest that the effect of chronic MHF feeding may extend beyond a simple reduction in hypothalamic odor responsiveness but rather reflects an altered sensory processing by selectively disrupting odor-evoked recruitment of ARC neurons. This possibility aligns with previous literature, showing that a chronic high-fat diet may disrupt synaptic processes within the ARC, including a reduction in excitatory postsynaptic current amplitude in ARC neurons (Wang and Li., 2022), suggesting that dietary exposure can remodel the synaptic properties of neurons involved in energy homeostasis. These changes in excitatory synaptic transmission could alter the strength of inputs received by ARC neurons and may contribute to the lack of odor-evoked activity observed following MHF feeding.

Moreover, the increased Iba-1 positive microglial labelling and close apposition of microglial processes to ODORTRAP labelled neurons in the ARC in DIO mice further suggests that changes in the hypothalamic environment may contribute to an altered neuronal response (Yi et al., 2013). Although the cellular identity and synaptic mechanisms underlying the reduced recruitment of odor-evoked ARC neurons remains undetermined, these findings suggest that chronic exposure to MHF may modify the sensory input recognition of hypothalamic feeding-related nuclei.

IPT exposure resulted in elevated NAc ODORTRAP activity in CF animals, while MHF animals displayed robust downregulation. Interestingly, ODORTRAP activity observed in our study was highly localized to the NAc medial shell subdivision, an area that receives dense innervation from lateral septum (Ma et al., 2020) and also classically from the VTA that exerts motivational salience by regulating glutamatergic transmission in the NAc medial shell (Yu et al., 2019). Additionally, AON innervates the NAc core and PC innervates the NAc lateral shell subdivision (Ma et al., 2020); a region where we do not observe any odor-induced TRAP activity. Hence, we speculate that the predominant inputs into the NAc medial shell subdivision are from either VTA or lateral septum. Given that we do not observe any differences in TRAP activity in the VTA of both CF and MHF animals in MO and IPT exposed groups, we speculate the non-involvement of these VTA – NAc medial shell inputs regulating the TRAP activity. Instead, the LS seems to be the likely candidate regulating NAc medial shell TRAP activation. Recent evidences have also demonstrated that M/TCs directly innervate the septum in mouse – in turn regulating their activity (Bulk et al., 2025). Hence the septum to NAc medial shell pathway might be crucial, engaging to assign motivational value to the odor stimulation (West and Carelli, 2016).

IPT exposure robustly activated MS neurons in CF animals that is in-line with recent report where chow odor stimulation elevated c-Fos and ps6 levels in MS glutamatergic neurons (Bulk et al., 2025). These neurons have been shown to play a role in anticipatory satiety signalling, so it is interesting that we observed that obese mice completely lost ODORTRAP activity in the MS. Our ODORTRAP data show strong septal labelling in control-fed animals but a marked reduction in MHF-fed mice; indicating an obesity-driven blunting of olfactory septal circuit activity.

We employed Iba-1 immunolabelling in the ARC of CF and MHF ODORTRAP animals to assess gliosis and neuroinflammation. Iba-1 is an established inflammatory marker validated in previous studies with pronounced increase in the ARC during DIO (Valdearcos et al., 2014). Moreover, the increased Iba-1 positive microglial labelling and close apposition of microglial processes to ODORTRAP labelled neurons in the ARC in DIO mice further suggests that changes in the hypothalamic environment may contribute to an altered neuronal response (Yi et al., 2013). Although the cellular identity and synaptic mechanisms underlying the reduced recruitment of odor-evoked ARC neurons remains undetermined, these findings suggest that chronic exposure to MHF may modify the sensory input recognition of hypothalamic feeding-related nuclei. Previous reports demonstrate the capacity of microglial activity within the ARC to directly modulate the activity of AgRP/NPY neurons (Reis et al., 2015). This supports our observation of Iba-1 cells in close apposition to isolated ODORTRAP neurons in the ARC of MHF animals potentially affecting their activity and feeding behaviour in DIO animals.

## 5. CONCLUSIONS

Together, our TRAP-based mapping reveals a discrete olfactory-hypothalamic-limbic network, with strong recruitment across olfactory cortex and feeding-related hypothalamic nodes. Importantly, our findings are further supported by prior work from our laboratory demonstrating that metabolic state robustly modulates the excitability of the olfactory bulb circuit and sensory processing. Earlier studies established that DIO and altered insulin signalling reshape olfactory processing and subsequent behavioral output (Fadool et al., 2004, 2011; Thiebaud et al., 2014; Chelette et al., 2022).

## Data Availability Statement

Data will be made available upon request from the corresponding author.

## Acknowledgements

This work was supported by grant R01DK133464 from the National Institutes of Health (NIDDK) and a VP for Research Postdoctoral Scholar Appointment from Florida State University to support grant T32DC000044 from the National Institutes of Health (NIDCD). We would like to thank Dr. Liz Brown for access to supplemental microscopy equipment in her laboratory and Dr. James M. Fadool for technical advice on the design/interpretation of our study.

## Abbreviations

glp1: glucagon-like peptide 1
M/TCs: mitral and tufted cells
OB: olfactory bulb
4-OHT: 4-hydroxytamoxifen
ADP: anterodorsal preoptic nucleus
AMY: amygdala
AON: anterior olfactory nucleus
ARC: arcuate nucleus
ARCa: anterior arcuate nucleus
ARCc: caudal arcuate nucleus
AVMA: American Veterinary Medicine Association
AVP: anteroventral preoptic nucleus
AVPV: anteroventral periventricular nucleus
BLA: basolateral amygdala
BS: blocking solution
CF: control fed
CoA: cortical amygdala
CreERT2: inducible modified cre recombinase
CRH: corticotropin releasing hormone
CTX: cortex
DAPI: 4′,6-diamidino-2-phenylindole
DIO: diet induced obesity
DMH: dorsomedial hypothalamus
DMHa: anterior dorsomedial hypothalamus
DMHc: caudal dorsomedial hypothalamus
DTT: dorsal tenia tecta
EPL: external plexiform layer
FSU: Florida State University
FosTRAP2 mice: B6;129-Fos2A-CreER
FOSTRAP mice: FosTRAP2 x Ai14
GC: granule cell
GCL: granule cell layer
glp1: glucagon-like peptide 1
GLM: glomerular layer
IACUC: Institutional Animal Care and Use Committee
ieg: immediate-early gene
IPT: isopropyl tiglate
i.p: intraperitoneal
LEC: lateral entorhinal cortex
LH: lateral hypothalamus
LHa: anterior lateral hypothalamus
LHc: caudal lateral hypothalamus
M/TCs: mitral and tufted cells
MC: mitral cell
MCH: melanin concentrating hormone
MCL: mitral cell layer
MHF: moderate high fat
MOB: main olfactory bulb
MO: mineral oil
MPN: medial preoptic nucleus
MS: medial septum
NAc: nucleus accumbens
Na HPO: disodium hydrogen phosphate
NaH PO: monosodium hydrogen phosphate
NGS: normal goat serum
NIH: National Institutes of Health
OB: olfactory bulb
OE: olfactory epithelium
Olfr160: olfactory receptor
OSN: olfactory sensory neurons
OT: olfactory tubercle
PB: phosphate buffer
PBS: phosphate-buffered saline
PFA: paraformaldehyde
PET: positron emission tomography
PM: permeabilization solution
POA: preoptic area
pPC: posterior piriform cortex
PVH: paraventricular hypothalamus
PVT: paraventricular nucleus of the thalamus
rt: room temperature
SCN: suprachiasmatic nucleus
SEM: standard error of the mean
SNc: substantia nigra pars compacta
TRAP: targeted recombination of active populations
TRAP2: Fos2A-iCreER mouse line
TRH: thyrotropin releasing hormone
VTA: ventral tegmental area
VMH: ventromedial hypothalamus
VTT: ventral tenia tecta

